# Temperature and intrinsic Ca^2+^ reshape TRPM4 pharmacology

**DOI:** 10.64898/2026.01.31.703029

**Authors:** Jinhong Hu, Sofia Ievleva, Sung Jin Park, Junuk Lee, Jennifer Cheng, Garrett O’Dea, Jiangnan Sheng, Juan Du, Wei Lü

## Abstract

Proteins operate in dynamic environments where ions, lipids, and temperature collectively define their properties, yet most studies rely on simplified conditions that overlook these intrinsic variables. Here, we show two such factors—temperature and Ca^2+^— remodel the function and pharmacology of TRPM4, an ion channel implicated in cardiac conduction, immune regulation, cancer, and intestinal fluid homeostasis. At physiological temperature and Ca^2+^, TPPO—previously considered a selective TRPM5 inhibitor inactive toward TRPM4—potently activates TRPM4, revealing strong synergy among temperature, Ca^2+^, and ligand binding. In contrast, Necrocide-1, a necroptotic activator targeting the same binding pocket, defies this logic: it opens TRPM4 without Ca^2+^ but is antagonized by Ca^2+^. Meanwhile, the inhibitors NBA and CBA engage a nearby pocket, locking the channel in a non-conductive pre-open state. Our findings highlight that even rigid binding pockets can exhibit temperature-dependent ligand recognition, revealing hidden pharmacology and informing selective, environment-aware therapeutic strategies.

## Introduction

Protein activity is inherently governed by the cellular environment in which it operates. A wide array of factors—including ions, lipids, temperature, pH, and protein–protein interactions—collectively determine how a protein functions in its native context. Yet, because of technical constraints, most biophysical and pharmacological studies have been conducted under simplified conditions, which have yielded foundational insights but often fail to capture the full physiological complexity.

Increasing efforts now aim to bridge this gap by incorporating more native variables—for example, using nanodiscs, liposomes or membrane-derived vesicles to preserve native lipidic environments for membrane proteins^1–4^, and purifying proteins from native sources to retain endogenous protein compositions and modifications^5–8^. Despite these advances, one fundamental determinant, temperature, is largely overlooked in mechanistic and pharmacological studies, even though it influences every protein in the human body, where proteins operate at 37°C. Because most in vitro assays are performed at room temperature or lower, therefore, the broad impact of temperature on protein conformation, ligand recognition, efficacy and mechanism remains unexplored, with only a few recent exceptions^9,10^.

Alongside temperature, intracellular Ca^2+^ represents another intrinsic physiological variable that exerts pervasive influence on protein behavior. Although not universal, Ca^2+^ acts as a selective yet broad regulator of cellular physiology. It is among the most tightly controlled cellular cofactors, with cytosolic concentrations fluctuating dynamically from tens of nanomolar at rest to low micromolar during signaling, and rising to abnormally high levels under pathological conditions such as ischemia, excitotoxicity, or cancer^11^. Many proteins are directly regulated by Ca^2+^ through dedicated binding sites, while others respond indirectly through Ca^2+^-dependent signaling cascades. Although the structural and functional roles of Ca^2+^ have been extensively characterized, its impact on protein–ligand interactions—particularly in combination with other intrinsic factors like temperature—has rarely been considered. This represents an important gap in our understanding of how Ca^2+^-regulated proteins engage ligands under physiological and pathological conditions.

The Ca^2+^-activated, temperature-sensitive channel Transient Receptor Potential Melastatin 4 (TRPM4) provides an ideal model to investigate how intrinsic physiological factors remodel pharmacology^12–15^. TRPM4 is a non-selective monovalent cation channel whose activation by elevated cytosolic Ca^2+^—arising from receptor-mediated signaling, Ca^2+^ entry through other channels, or release from intracellular stores—leads to Na^+^ influx and membrane depolarization. This depolarization activates downstream depolarization-dependent mechanisms, including the regulation of voltage-gated ion channels, and thereby modulates diverse downstream processes such as cardiac conduction, insulin secretion, neuronal excitability, immune regulation, and intestinal fluid homeostasis^16–25^. Pathogenic variants have been implicated in Brugada syndrome, a genetic heart rhythm disorder, and other inherited conduction disorders, as well as diverse cancers^26–34^. In this context, TRPM4 antagonists may be therapeutically beneficial in cardiac diseases where excessive TRPM4 activity contributes to pathological depolarization and conduction defects, whereas TRPM4 agonists may be relevant in certain cancers in which TRPM4 is upregulated, where enhanced channel activity has been associated with necrotic cell death pathways. Despite this broad physiological and pathological relevance, TRPM4 has remained largely untargeted by clinical pharmacology, with the single exception of bisacodyl—a widely used drug for chronic constipation—and its active metabolite deacetyl bisacodyl, which we recently identified and characterized as clinically relevant TRPM4 agonists^25^.

Our previous structural and functional studies revealed that TRPM4 adopts two major conformations, a “cold” and a “warm” state, whose equilibrium is governed jointly by temperature and cytosolic Ca^2+^ concentration^10^. In the cold conformation, Ca^2+^ binds exclusively to an agonist site within the S1–S4 bundle (Ca_TMD_), giving rise to strongly outwardly rectifying currents. In contrast, in the warm conformation, the intracellular domain undergoes a pronounced rearrangement accompanied by Ca^2+^ binding at an additional intracellular site (Ca_warm_ or Ca_ICD_), resulting in currents that are less outwardly rectifying and exhibit substantial inward conductance. Together, these findings suggest that biophysical and pharmacological assays performed under room temperature and undefined Ca^2+^ conditions may not accurately capture the channel’s behavior under physiological settings.

Here, we systematically examine TRPM4 pharmacology under conditions that incorporate both temperature and Ca^2+^. Through combined cryo-EM and electrophysiology, we reveal that these intrinsic factors profoundly shape ligand recognition, efficacy, and mechanism. Our results identify the S1–S4 domain as a dynamic regulatory hub that integrates environmental and chemical cues to govern activation and inhibition. More broadly, this work establishes an environment-aware framework for pharmacology, in which incorporating intrinsic variables exposes hidden drug activities and new mechanisms of regulation. For widely expressed proteins like TRPM4, such principles open opportunities to design therapeutics that exploit pathological local conditions—such as abnormally elevated Ca^2+^ or altered thermal states—for selective action in disease, while preserving normal physiological function and minimizing side effects.

## Results

### Temperature reveals hidden pharmacology of TRPM4

During the cold-to-warm transition of TRPM4, Ca^2+^ and temperature drive major conformational changes in the intracellular domain but only subtle shifts in the transmembrane domain^10^. From these structures, we identified three categories of ligand binding sites that emerge elusively in the warm conformation of the intracellular domain^10^. While this outcome may be anticipated from the magnitude of ICD rearrangement, it nevertheless raises two central questions: whether drug screening performed exclusively at room temperature may have overlooked active compounds, and whether temperature-dependent ligand recognition is confined to regions undergoing large conformational change (such as the intracellular domain) or extends to regions with more modest movements (such as the transmembrane domain).

To address these questions, we examined several previously reported TRPM4 ligands, as well as compounds considered inactive toward TRPM4. We focused on hydrophobic ligands predicted to interact with the transmembrane domain, including triphenylphosphine oxide (TPPO), necrocide-1 (NC1), and the anthranilic acid derivatives CBA (4-chloro-2-[2-(2-chloro-phenoxy)-acetylamino]-benzoic acid) and NBA (4-chloro-2-(2-(naphthalene-1-yloxy) acetamido) benzoic acid). TPPO, originally identified as a selective TRPM5 inhibitor with an IC_50_ of 12 μM, has been found inactive on TRPM4 when tested using fluorescence-based membrane potential assays^35^. NC1 was recently characterized as a TRPM4 activator that promotes Na^+^ influx and triggers necrotic cell death^36^, suggesting therapeutic potential in cancer. NBA and CBA represent the best-characterized TRPM4 inhibitors to date with IC_50_ in the sub-micromolar range^37^. Importantly, all prior functional and structural studies were conducted at room temperature or lower.

Using patch-clamp electrophysiology, we tested these compounds at both room temperature and 37°C. Under previously reported assay conditions—room temperature with either zero free intracellular Ca^2+^ or basal free intracellular Ca^2+^ (100 nM), we reproduced established results: 50 μM TPPO showed negligible activity on TRPM4—comparable to the control group, where basal Ca^2+^ alone is insufficient to drive channel activation (Fig. 1a,b; Extended Data Fig. 1a,b); NC1 activated the channel, eliciting largely linear, voltage-independent currents (Fig. 2a,b); and NBA and CBA produced robust inhibition of Ca^2+^-induced TRPM4 current (Extended Data Fig. 2a).

**Figure 1.**
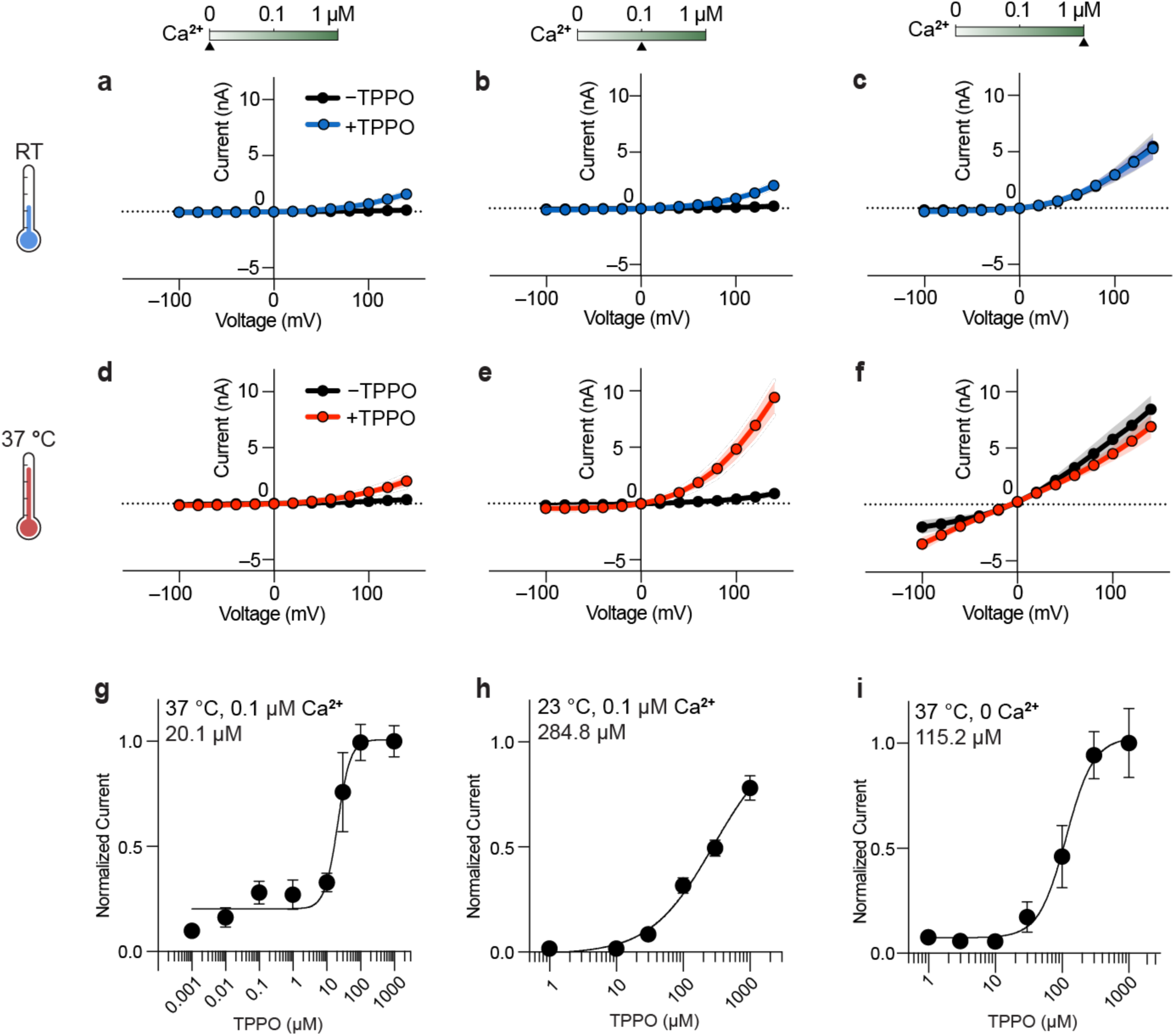
Three-way synergy between TPPO, temperature and Ca^2+^. **a–f**, Whole-cell currents measured in tsA cells overexpressing wild-type TRPM4 at 22 °C (upper panels) and 37 °C (lower panels), with free intracellular Ca^2+^ concentrations of 0 µM (**a, d**), 0.1 µM (**b, e**) and 1 µM (**c, f**), in the absence or presence of 50 µM extracellular TPPO. A voltage protocol stepping from −100 mV to +140 mV in 20-mV increments was applied, with each voltage step lasting 100 ms. The currents from all measured cells were then averaged and plotted versus voltage to generate the mean I–V curve, with the line representing the mean and the shaded indicating the s.e.m. The numbers of independently measured cells (n) are as follows: (**a**) n=9 (−TPPO) / 9 (+TPPO), (**b**) n=7 (−TPPO) / 8 (+TPPO), (**c**) n=9 (−TPPO) / 9 (+TPPO), (**d**) n=7 (−TPPO) / 10 (+TPPO), (**e**) n=8 (−TPPO) / 8 (+TPPO), and (**f**) n=6 (−TPPO) / 6 (+TPPO). **g–i**, TPPO dose-response measurements for wild-type TRPM4 with 0.1 µM free intracellular Ca^2+^ at 37 °C (**g**), 0.1 µM Ca^2+^ at 23 °C (**h**), and 0 µM Ca^2+^ at 37 °C (**i**). A voltage protocol was applied every 10 s to monitor current changes: –100 mV for 200 ms, followed by +100 mV for 200 ms. Once the current stabilized at each TPPO concentration, the current amplitudes at +100 mV from all measured cells were averaged, and the mean amplitudes were plotted and fitted to obtain the EC_50_. Current amplitudes were normalized either to the mean at 1000 µM TPPO (**g, i**) or to the fitted I_max from the Hill equation (**h**). Data are presented as mean ± s.e.m. The numbers of independently measured cell (n) are as follows: (**g**) n=7 for all concentrations, (**h**) n=5 at 1 µM and n=7 for all other concentrations, and (**i**) n=7 for all concentrations.

**Figure 2.**
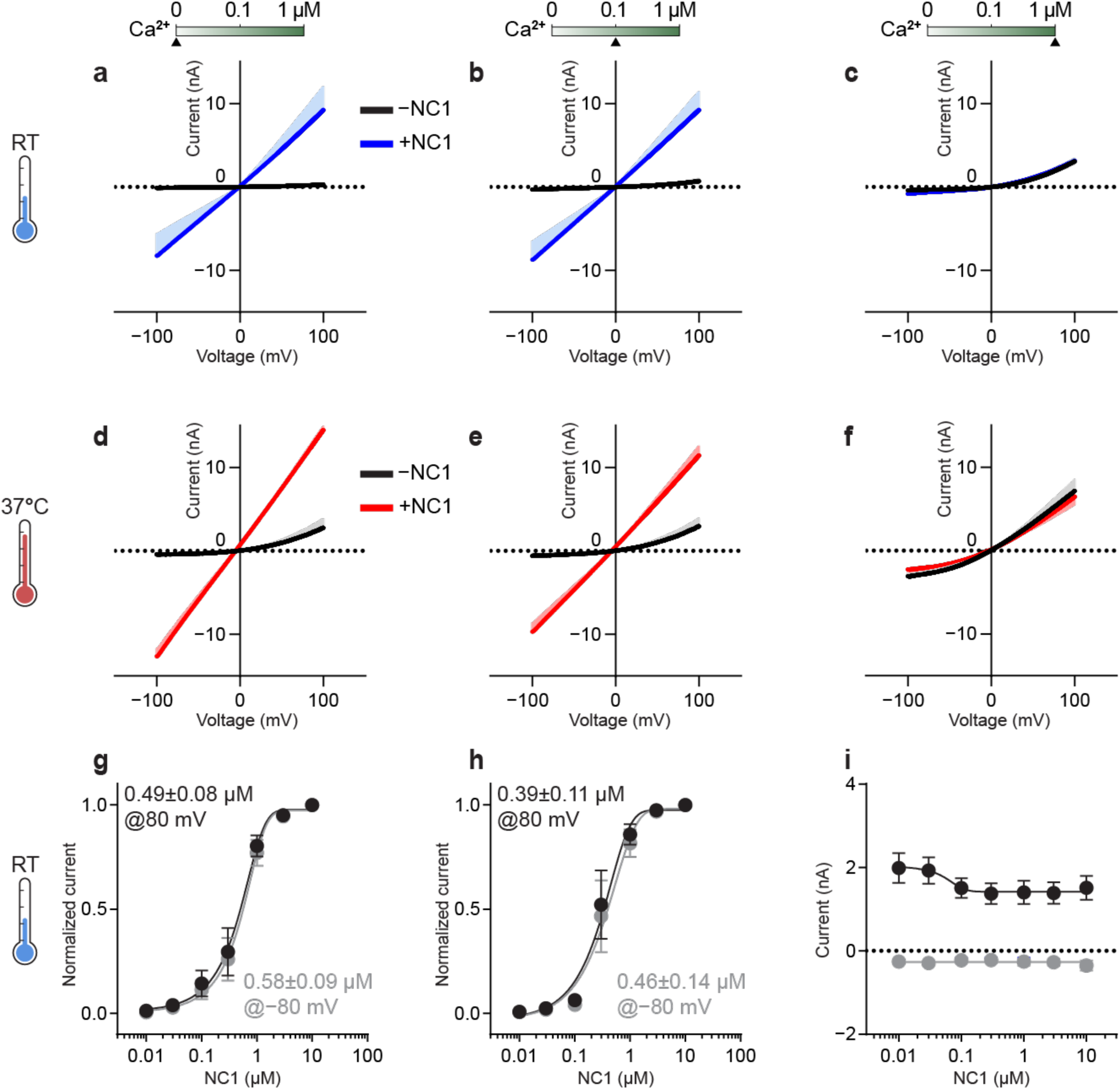
Ca^2+^-dependent gating reversal of NC1. **a–f**, Whole-cell currents measured in tsA cells overexpressing wild-type TRPM4 at 22 °C (upper panels) and 37 °C (lower panels), with free intracellular Ca^2+^ concentrations of 0 µM (**a, d**), 0.1 µM (**b, e**) and 1 µM (**c, f**). A voltage protocol was applied every 5 s to monitor current changes until reaching steady state: −100 mV for 50 ms, ramped to +100 mV over 200 ms and held at +100 mV for 50 ms. The currents without and with 1 µM extracellular NC1 in each condition were measured in a single cell. The currents from all measured cells were then averaged and plotted versus voltage to generate the mean I–V curve, with the line representing the mean and the shaded indicating the s.e.m. For clarity, the s.e.m. envelope was plotted in one direction. The numbers of independently measured cells (n) are as follows: n=5 (**a**), n=7 (**b**), n=5 (**c**), n=5 (**d**), n=4 (**e**), and n=4 (**f**). **g–i**, NC1 dose-response measurements for wild-type TRPM4 at 22 °C with 0 µM free intracellular Ca^2+^ (**g**), 0.1 µM Ca^2+^ (**h**), and 1 µM Ca^2+^ (**i**). A voltage protocol was applied every 5 s to monitor current changes: 0 mV for 50 ms, switch to +80 mV for 50 ms, followed by −80 mV for 50 ms, and return to 0 mV for 50 ms. Once the current stabilized at each NC1 concentration, the current amplitudes at +80 mV and −80 mV were normalized to those at 10 µM NC1. Normalized currents from all measured cells were averaged and plotted as mean ± s.e.m. EC_50_ values obtained from individual cells were averaged to yield the final EC_50_ reported as mean ± s.e.m. (**g**,**h**). Data are presented as mean ± s.e.m. The numbers of independently measured cells (n) are as follows: n=5 (**g**), n=5 (**h**), and n=4 (**i**).

Strikingly, elevating the temperature to 37 °C revealed a hidden pharmacological behavior. While NC1, NBA, and CBA retained their room-temperature profiles (Fig. 2d,e; Extended Data Fig. 2a), TPPO displayed a completely different response—robustly activating TRPM4 currents (Fig. 1e; Extended Data Fig. 1e). These observations highlight a critical gap in conventional pharmacological screening: assays performed under non-physiological conditions may both overlook potential active compounds and misclassify ligand selectivity.

### Ca^2+^ as a key determinant of agonist recognition and potency in TRPM4

Ca^2+^ is the only known endogenous agonist of TRPM4, and together with temperature, drives the transition between its cold to warm conformations^10^. Building on our findings that temperature reshapes TRPM4 pharmacology (Fig. 1e), we next explored whether Ca^2+^ modulates the binding and potency of the exogenous activators TPPO and NC1.

Under Ca^2+^-free conditions achieved by EGTA buffering, 50 uM TPPO produced negligible activation of TRPM4 (Fig. 1d; Extended Data Fig. 1d), whereas adding 100 nM free intracellular Ca^2+^ enabled robust activation (Fig. 1e, red traces; Extended Data Fig. 1e). Thus, even basal Ca^2+^ concentrations—although insufficient to open the channel alone (Fig. 1e, black trace; Extended Data Fig. 1e)—markedly enhance TPPO-induced TRPM4 activation. To quantify this apparent synergy, we determined TPPO dose-response relationships. Physiological temperature increased TPPO potency by ∼14-fold under basal Ca^2+^ (Fig. 1g,h; Extended Data Fig. 1g,h), while basal Ca^2+^ increased TPPO potency by ∼6-fold at 37 °C (Fig. 1g,i; Extended Data Fig. 1g,i). In the absence of both Ca^2+^ and physiological temperature, TPPO activated TRPM4 only weakly at submillimolar concentrations and failed to reach saturation even at millimolar levels. When intracellular Ca^2+^ reach activating level (1 µM), the effect of TPPO is largely—though not completely—masked. Under these conditions, the current recorded in the presence of TPPO appeared relatively linear and closely resembled that observed in the absence of TPPO, a phenotype we previously reported for TRPM4 activated by 1 µM Ca^2+^ at 37 °C^10^ , aside from minor differences that may reflect a combined contribution of TPPO and Ca^2+^. Together, these observations are consistent with TPPO and Ca^2+^ acting through a shared activation pathway, with Ca^2+^ becoming the dominant determinant of the channel behavior at high concentrations (Fig. 1c,f; Extended Data Fig 1c,f).

Together, these data demonstrate that physiological context is essential for TPPO efficacy, revealing a three-way synergy between temperature, Ca^2+^, and ligand binding. This context dependence overturns the view that TPPO is inactive toward TRPM4, revealing instead that it potently activates the channel—opposite to its inhibitory action on the closely related TRPM5^35^. This finding underscores how ligand behavior can be fundamentally mischaracterized when assays are performed under non-physiological conditions.

We next examined whether Ca^2+^ influences NC1, previously reported as the first Ca^2+^-independent agonist of TRPM4^36^. Under Ca^2+^-free and 100 nM Ca^2+^ conditions, NC1 evoked nearly identical currents with linear current-voltage relationships distinct from Ca^2+^-activated currents, confirming that NC1 can activate TRPM4 independently with an EC_50_ in the submicromolar range (Fig. 2a,b,d,e,g,h; Extended Data Fig. 3a). However, at 1 µM Ca^2+^, a physiologically relevant level during cellular stress that robustly activates TRPM4, the response changed dramatically, becoming strongly outwardly rectified (Fig. 2c,f)—a hallmark of canonical Ca^2+^-mediated activation^14^. This striking reversal suggests that NC1 is not simply Ca^2+^-independent: once Ca^2+^ reaches activating levels, its action dominates, functionally masking NC1-mediated gating even at saturating NC1 concentrations (Fig. 2i). Thus, rather than bypassing the Ca^2+^ requirement, NC1 displays a biphasic Ca^2+^ dependence: permissive at basal Ca^2+^, yet exhibiting little to no additional effect at higher Ca^2+^ concentrations. This behavior is most consistent with NC1 becoming functionally masked or antagonized at elevated Ca^2+^, either through reduced/abolished NC1 binding due to Ca^2+^-dependent competition, or through allosteric coupling between NC1- and Ca^2+^-binding sites in scenarios where co-binding occurs. Which mechanism is more likely is discussed further in the section “Mechanistic basis for Ca^2+^-dependent gating reversal of NC1”.

Taken together, these results reveal distinct modes of Ca^2+^-agonist interplay in TRPM4. TPPO requires basal Ca^2+^ (and temperature) to act effectively on the channel, whereas NC1 functions independent of Ca^2+^ at rest but loses efficacy once Ca^2+^ rises to stimulatory levels. This contrast illustrates how the same intrinsic cofactor can differentially tune ligand action through cooperative or antagonistic coupling. More broadly, they reinforce the importance of evaluating ion channel pharmacology under physiological conditions, as environmental context can fundamentally alter both efficacy and apparent selectivity.

### Structural basis of the three-way synergy between TPPO, temperature and Ca^2+^

To elucidate how temperature and Ca^2+^ shape TPPO activation of TRPM4, we determined cryo-EM structures of the channel in the presence of TPPO at 18 °C and 37 °C, under both Ca^2+^-free and Ca^2+^-saturated conditions. All the structures were resolved to high quality at ∼2.5–2.9 Å resolutions, allowing unambiguous identification of ligand binding sites (Extended Data Figs. 4–5; Supplementary Fig. 1; Supplementary Data Table 1).

In the consensus map of the Ca^2+^/TPPO–TRPM4–37°C dataset, we observed a prominent tripod-shaped density in the S1–S4 domain of each protomer. This feature closely matches the shape of TPPO, and sits immediately above the Ca_TMD_ site (Fig. 3a). We therefore designated this pocket as the S1–S4_Upper_ site. This site spatially overlaps with the deacetyl bisacodyl binding pocket we identified previously^25^. Subunit-level classification revealed that where warm and cold conformations coexist in this dataset, with the warm state predominating (Extended Data Fig. 4). The ligand density is markedly stronger in the warm state, whereas a weaker yet still discernible density exists in the cold state (Fig. 3d; this minor cold state is discussed at the end of this section). Within this pocket, TPPO occupies a hydrophobic cavity and forms a critical polar interaction between its phosphoryl moiety and R1072 on the TRP helix (Fig. 3d, left panel). To validate this assignment, we mutated key coordinating residues to alanine and performed electrophysiological recording. Most mutations impaired Ca^2+^-dependent activation, likely by disrupting the integrity of the adjacent Ca_TMD_ site, making it impossible to unambiguously isolate their effects on TPPO, as TPPO requires Ca_TMD_ to manifest its activity (Extended Data Fig. 1j).

**Figure 3.**
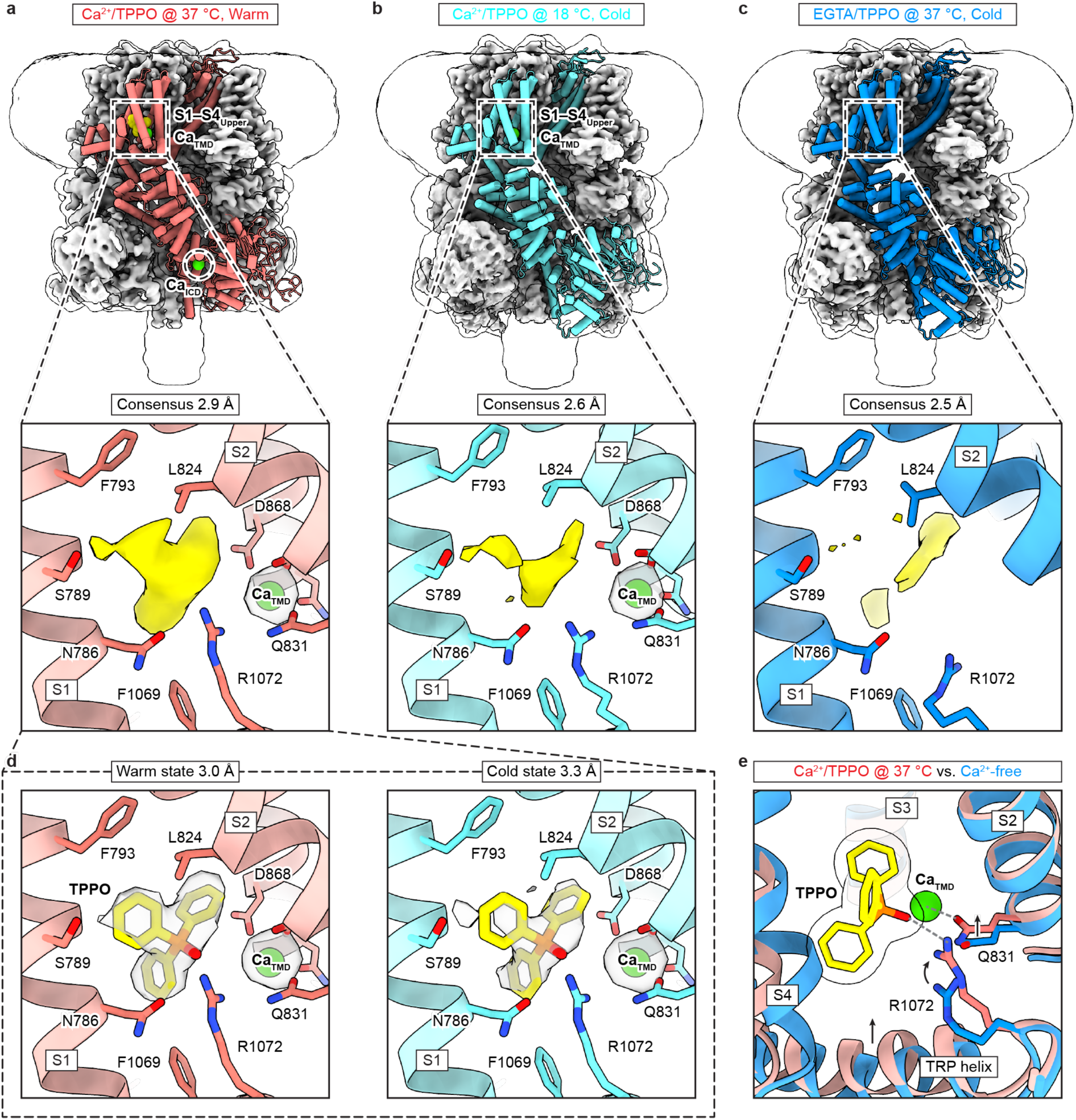
Structural basis of the three-way synergy between TPPO, temperature and Ca^2+^. **a–c**, Structures of TRPM4 in complex with Ca^2+^/TPPO at 37 °C (**a**, PDB-ID 9Z1W), Ca^2+^/TPPO at 18 °C (**b**, PDB-ID 9Z1Z), and EGTA/TPPO at 37 °C (**c**, PDB-ID 9Z20). One subunit is shown in cartoon representation, while the remaining three subunits are displayed as the cryo-EM map, viewed parallel to the membrane. The transparent envelope outlines the detergent micelle and the disordered region of the C-terminal coiled-coil. TPPO is highlighted in yellow and Ca^2+^ in green. The resolution of the consensus map is indicated below. Bottom panels show the details of the S1–S4 binding site in the consensus maps. The protein backbone is shown in cartoon representation, with key interacting residues in stick representation. Densities at the TPPO binding site (yellow) and at the Ca_TMD_ site (transparent in **a**,**b**) are contoured at 4.5 σ. **d**, Details of the S1–S4 binding site in the warm (left, PDB-ID 9Z1X) and cold (right, PDB-ID 9Z1Y) states, obtained from classification of the Ca^2+^/TPPO dataset collected at 37 °C. The resolution of each cryo-EM map is indicated. Densities at the TPPO binding site and the Ca_TMD_ site are shown as transparent surface representations contoured at 9 σ. The fitted TPPO molecule is shown in stick representation, and Ca_TMD_ as a green sphere. **e**, Structural comparison of TRPM4 with Ca^2+^/TPPO at 37 °C (salmon, PDB-ID 9Z1X) and the Ca^2+^-free apo state (blue, PDB-ID 9B93), superimposed using the S1–S4 (residues 760–910). TPPO is shown in stick representation with a transparent envelope, and Ca_TMD_ as a green sphere. Black arrows indicate the approximate direction of movement of key residues and the TRP helix. The dashed lines mark key interactions.

In contrast, R1072A preserved Ca^2+^-evoked gating yet abolished TPPO-dependent activation, thereby confirming the functional relevance of the TPPO binding site (Extended Data Fig. 1k).

In the Ca^2+^-bound cold conformation from the Ca^2+^/TPPO–TRPM4–18°C dataset, only a faint and incomplete signal was discernible at the S1–S4_Upper_ site at very low contour levels (Fig. 3b). The density formed a triangular outline reminiscent of TPPO but was markedly smaller than the full ligand, consistent with partial or unstable occupancy. In the Ca^2+^-free cold conformation from the EGTA/TPPO–TRPM4–37°C dataset, this feature disappeared entirely, with no trace of TPPO density detectable (Fig. 3c).

Thus, TPPO binding exhibits a clear dependence on the cold–warm conformational equilibrium of TRPM4, a transition that is cooperatively governed by temperature and Ca^2+^ (ref^10^). This structural observation coincides with our electrophysiological finding that TPPO, temperature, and Ca^2+^ act in three-way synergy to promote channel activation (Fig. 1a,b,d,e). Together, these results suggest that TPPO efficacy is tightly coupled to the conformational landscape defined by these two intrinsic modulators.

To dissect the mechanism underlying this coupling, we compared the Ca^2+^-free cold and Ca^2+^/TPPO-bound warm structures. In the Ca^2+^-free cold conformation, R1072 is displaced from the S1–S4_Upper_ pocket and stabilized by an interaction with Q831 on S2—a key residue involved in Ca_TMD_ binding—thereby precluding TPPO binding (Fig. 3e). Upon transition to the warm state, upward movement of the intracellular domain tilts the TRP helix, moving R1072 into position for TPPO coordination (Fig. 3e). Concurrently, Ca^2+^ binding at the Ca_TMD_ site recruits Q831 for Ca^2+^ coordination, releasing R1072 from its locked pose for direct interaction with the TPPO phosphoryl group (Fig. 3e). Together, these coupled rearrangements prime the S1–S4_Upper_ pocket for effective ligand engagement.

Interestingly, a weak TPPO-shaped density was also detected in the minor Ca^2+^-bound cold conformation present in the Ca^2+^/TPPO–TRPM4–37°C dataset, identified through single-subunit classification (Extended Data Fig. 4). This density was weaker than in the warm conformation but stronger than in the corresponding Ca^2+^-bound cold conformation resolved at 18°C (Fig. 3b, lower panel; Fig. 3d; Extended Data Fig. 6), indicating that TPPO occupancy increases with temperature even within the same overall protein conformation. Because the backbone architecture of the binding site remains nearly identical between the two cold structures, this enhancement likely arises from subtle, temperature-dependent sidechain rearrangements—below the resolution of our maps (∼2.6 Å local)—that fine-tune ligand packing and render the interaction thermodynamically more favorable.

Together, these results reveal that TPPO selectively engages the warm conformation of TRPM4 by exploiting temperature- and Ca^2+^-dependent changes in local binding energetics. This provides a direct structural basis for the observed three-way synergy, illustrating how intrinsic modulators reshape the binding pocket to control ligand efficacy at the atomic level.

### Mechanistic basis for Ca^2+^-dependent gating reversal of NC1

Building on our observation that NC1 activates TRPM4 in the absence of Ca^2+^ but loses efficacy once Ca^2+^ reaches activating levels (Fig. 2a–f), we sought to define the structural basis of this gating reversal. To this end, we determined cryo-EM structures of TRPM4 in complex with NC1 at 37 °C under Ca^2+^-free and Ca^2+^-bound conditions, resolved to 2.6 and 2.8 Å, respectively (Extended Data Figs. 7; Supplementary Fig. 1; Supplementary Data Table 2).

In the Ca^2+^-free dataset, a well-defined tripod-shaped density was observed in the S1–S4 domain (∼2.6 Å local resolution), occupying the same pocket as the TPPO and deacetyl bisacodyl binding sites^25^, while no Ca^2+^ density appeared at the Ca_TMD_ site, as expected (Fig. 4a). The shape and dimensions of this density unambiguously matched NC1, which, despite being larger than TPPO, adopts a comparable three-ring scaffold with its indolinone carbonyl oxygen positioned analogously to TPPO’s phosphoryl oxygen (Figs. 3d, left panel; Fig. 4a).

**Figure 4.**
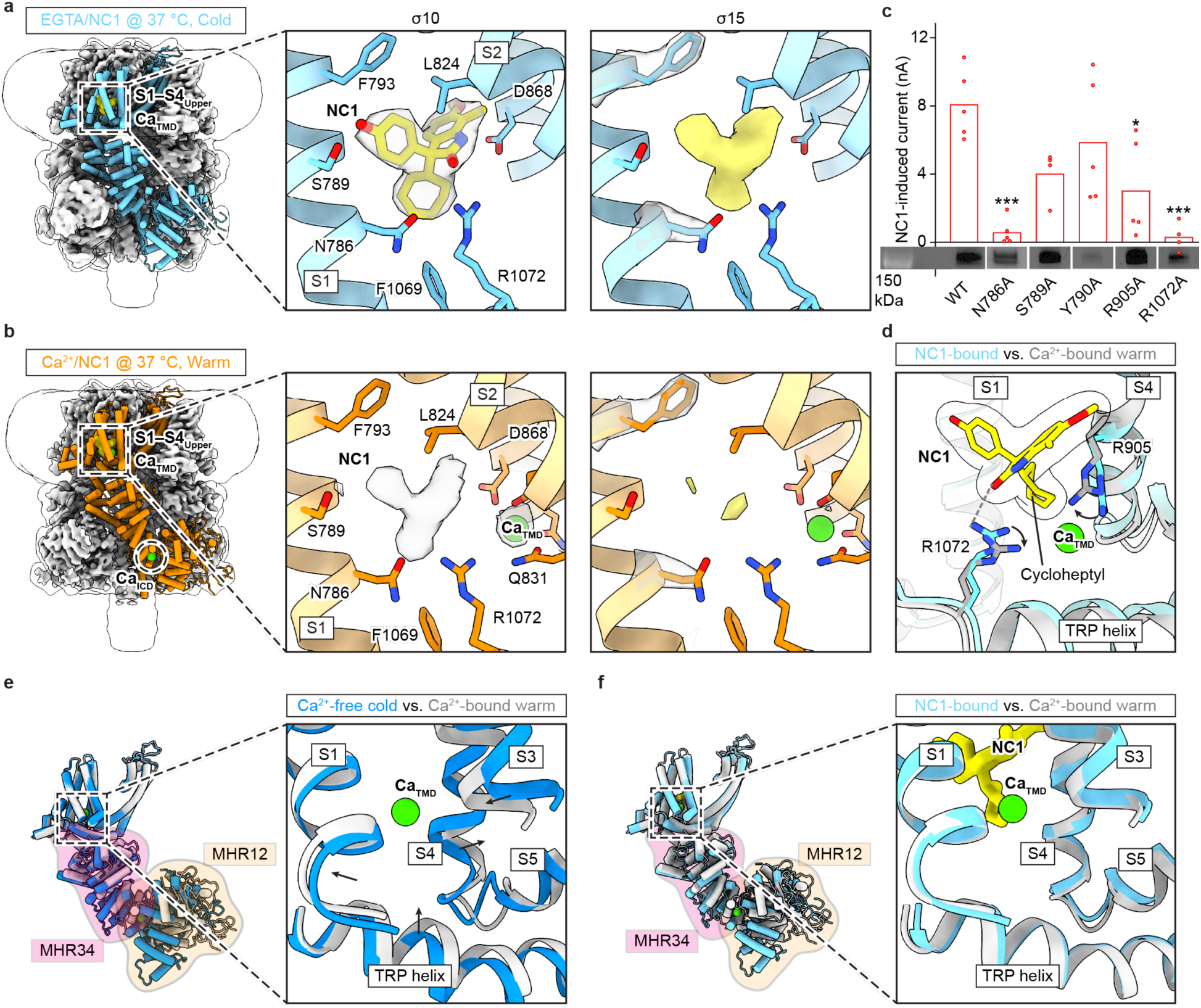
Mechanistic basis for Ca^2+^-dependent gating reversal of NC1. **a**,**b** The left panels show the structures of TRPM4 in complex with EGTA/NC1 at 37 °C (**a**, PDB-ID 9Z22) and Ca^2+^/NC1 at 37 °C (**b**, PDB-ID 9Z21). One subunit is shown in cartoon representation, while the remaining three subunits are displayed as the cryo-EM map, viewed parallel to the membrane. The transparent envelope outlines the detergent micelle and the disordered region of the C-terminal coiled-coil. The middle panels show the details of the S1–S4 binding site, with the protein backbone in cartoon representation and key interacting residues in stick representation. Densities at the NC1 binding site, and at the Ca_TMD_ site (only in **b**) are shown as transparent surface representations contoured at 10 σ. The fitted NC1 molecule in (only in **a**) is shown in stick representation, and Ca_TMD_ as a green sphere (only in **b**). The right panels present the same regions as the middle panels but additionally display densities for well-defined side chains within the binding site, contoured at 15 σ. **c**, NC1-induced whole-cell current amplitudes at +100 mV in cells overexpressing wild-type TRPM4 and binding-site mutants, calculated by subtracting the steady-state current recorded before NC1 application from that recorded after application. Each point represents an independently measured cell, with bars indicating the mean. The numbers of independently measured cells are 5 (WT), 5 (N786A), 4 (S789A), 5 (Y790A), 5 (R905A), and 4 (R1072A). Statistical analysis was performed using one-way ANOVA with Bonferroni’s post hoc test, comparing each mutant with wild type (*p<0.05, ***p<0.001). The adjusted p-values for each mutant compared to the wild-type (WT) control were 0.0002 for N786A, 0.0741 for S789A, 0.7002 for Y790A, 0.0103 for R905A and 0.0002 for R1072A. Averaged raw data are shown in Extended Data Fig. 3b. Cell surface expression of wild-type TRPM4 and mutants is shown, confirming proper trafficking to the plasma membrane; uncropped gels are provided in Supplementary Fig. 2. **d**, Structural comparison of TRPM4 NC1-bound state (cyan, PDB-ID 9Z22) and the Ca^2+^-bound warm state (blue, PDB-ID 9B8W) of TRPM4, superimposed using the S1–S4 (residues 760–910). NC1 is shown in stick representation with a transparent envelope, and Ca_TMD_ as a green sphere. Black arrows indicate the approximate direction of movement of key residues. The dashed line marks a key interaction. **E**, Structural comparison of a single subunit of the Ca^2+^-free cold state (blue, PDB-ID 9B93) and the Ca^2+^-bound warm state (grey, PDB-ID 9B8W) of TRPM4, superimposed using the S1–S4 (residues 760–910). MHR3/4 is outlined with a pink envelope and MHR1/2 with an orange envelope. The right panel shows details of the S1–S4 binding site and its surroundings. Ca_TMD_ is shown in a green sphere, and the black arrows indicate the movement of structural elements forming the binding site upon Ca^2+^ binding. **F**, Structural comparison of the NC1-bound state (cyan, PDB-ID 9Z22) and the Ca^2+^-bound warm state (grey, PDB-ID 9B8W) of TRPM4. The structures are superimposed and displayed similarly to panel (**e**). NC1 is shown in stick representation, and Ca_TMD_ as a green sphere.

Consequently, NC1 binds in a similar pose to TPPO, engaging an equivalent network of residues, including a key polar contact between its indolinone carbonyl oxygen and R1072 on the TRP helix (Fig. 4a). Consistent with this, alanine substitution of R1072 and other residues within the binding site abolished or markedly reduced NC1-mediated activation (Fig. 4c; Extended Data Fig. 3b), confirming that NC1, like TPPO, binds to the S1–S4_Upper_ site.

Interestingly, in the Ca^2+^-bound dataset, NC1 density at the same S1–S4_Upper_ pocket was markedly weaker (Fig. 4b). This diminished occupancy aligns with our electrophysiological findings, where 1 uM Ca^2+^ strongly suppressed the linear, NC1-evoked currents characteristic of Ca^2+^-free conditions and restored the canonical Ca^2+^-dependent gating pattern (Fig. 2a–f).

Together, these data indicate that NC1 binding is strongly disfavored in the presence of Ca^2+^. While we cannot exclude the possibility that NC1 and Ca^2+^ may co-bind and produce an electrophysiological profile similar to Ca^2+^ alone, the pronounced weakening of NC1 density in the Ca^2+^-bound structures suggests that a Ca^2+^-dependent reduction, or near abolition, of NC1 binding is the most likely explanation. Accordingly, we interpret the weak residual density as arising from the oversaturating NC1 concentration used for cryo-EM rather than from stable co-binding. Importantly, the antagonism does not result from direct competition for a shared site but instead from allosteric interference between the NC1 pocket and the Ca_TMD_ site (Fig. 4b).

To understand the basis of this interference, we compared the Ca^2+^-free NC1–TRPM4 and Ca^2+^–TRPM4 complexes. Two local structural changes accompany Ca^2+^ coordination at the Ca_TMD_. First, Ca^2+^ binding repositions the TRP helix, shifting R1072 away from the indolinone carbonyl oxygen that is central to NC1 recognition (Fig. 4d). Second, Ca^2+^ binding brings the positively charged R905 into close proximity with NC1’s hydrophobic seven-membered cycloheptyl ring, creating an electrostatically unfavorable juxtaposition that further destabilizes ligand binding (Fig. 4d). Together, these antagonistic rearrangements explain why NC1 affinity drops at increased Ca^2+^ concentration.

In summary, NC1 represents the first example of a TRPM4 activator that is temperature-independent yet negatively regulated by intrinsic Ca^2+^. Structural and functional analyses together uncover a Ca^2+^-dependent gating reversal, whereby Ca^2+^ reshapes the allosteric network to displace a previously bound activator. This mechanism contrasts sharply with TPPO, whose efficacy requires Ca^2+^ synergy at physiological temperature, underscoring the opposing logic by which intrinsic modulators shape TRPM4 pharmacology.

### NC1 overrides Ca^2+^-dependent ICD–TMD allosteric regulation

The defining feature of NC1 is that it activates TRPM4 without requiring Ca^2+^ at either of the two regulatory sites (Fig. 2a,d): the agonist site Ca_TMD_ and the temperature-dependent allosteric site in the intracellular domain (Ca_ICD_). In canonical gating, Ca^2+^ and temperature cooperatively couple the ICD and TMD to drive activation: Ca^2+^ binding at the temperature-sensitive Ca_ICD_ shifts the channel from the cold to the warm state, establishing the primed conformation in which Ca^2+^ binding at the Ca_TMD_ opens the pore at physiologically relevant membrane potentials^10^. To understand how NC1 circumvents this mechanism, we compared three structural states: Ca^2+^-free cold, Ca^2+^-bound warm, and NC1-bound. The transition from Ca^2+^-free cold to Ca^2+^-bound warm involves major ICD rearrangements coupled to a substantial shift in the TRP helix and TMD (Fig. 4e). By contrast, in the NC1-bound structure the ICD remains in the cold-like, yet the TMD closely resembles the warm state conformation even in the absence of Ca^2+^ at the Ca_TMD_ (Fig. 4f). These observations suggest that NC1 functionally substitutes for Ca^2+^ at the TMD while operating independently of Ca^2+^-dependent priming at the ICD.

Electrophysiology supported this model: NC1 elicited robust currents of comparable amplitude at both room and physiological temperatures (Fig. 2b,e)—in contrast to Ca^2+^-dependent TRPM4 activation, which is strongly enhanced at 37 °C through Ca_ICD_-mediated ICD–TMD coupling^10,15^. Thus, NC1 activates the channel without engaging the Ca_ICD_-driven allosteric control. To directly test this mechanism, we examined the E396A mutant, which abolishes Ca_ICD_ binding and locks TRPM4 in the cold-like conformation regardless of temperature^10^. Remarkably, NC1 still produced strong, temperature-independent activation in this mutant, confirming that NC1 does not require the ICD–TMD regulation (Extended Data Fig. 3c). This unprecedented mode of action reveals an alternative allosteric pathway within TRPM4 and highlights NC1 as a promising scaffold for therapeutic development, particularly under conditions where intracellular Ca^2+^ fluctuates.

### NBA and CBA inhibit TRPM4 through the S1–S4_Lower_ site

Pharmacological inhibition of TRPM4 is of broad clinical importance, as channel overactivation contributes to arrhythmogenic cardiac disorders, ischemic injury, and cancer progression^18,21,38^. However, our previous work revealed that the endogenous antagonist ATP markedly loses potency at physiological temperature because its binding location shifts between the cold and warm conformations of the channel^10^. This temperature sensitivity limits ATP’s ability to suppress TRPM4 under normal cellular conditions^39^.

To identify inhibitors that remain effective within the physiological conformational landscape, we turned to NBA and its analog CBA^37^. In contrast to ATP and other temperature-sensitive ligands, NBA and CBA inhibit TRPM4 in a temperature-independent manner, maintaining high potency at 37 °C (Extended Data Fig. 2a). This makes them ideal probes for uncovering how TRPM4 inhibition can be achieved under physiological conditions through a temperature-insensitive mechanism.

At 37 °C in the presence of Ca^2+^, cryo-EM analysis of TRPM4 in complex with NBA or CBA uncovered a previously unknown ligand binding site within the S1–S4 domain in both datasets, present in both cold and warm conformations (Extended Data Figs. 8; Supplementary Fig. 1).

This site lies below the Ca_TMD_ site and adjacent to the S1–S4_Upper_ pocket that accommodates TPPO and NC1, and is therefore termed S1–S4_Lower_ site (Fig. 5a,b). The ligand densities were of excellent quality (local resolution ∼2.6 Å) and closely matched the crystal structures of NBA and CBA^40^, respectively, allowing unambiguous ligand placement (Supplementary Fig. 1; Extended Data Fig. 9a,b; Supplementary Tables 2, 3). In both inhibitors, the chlorophenyl-carboxyl group points upward within the S1–S4 domain, with its carbonyl stabilized by two positively charged residues, R905 on S4 and R1072 on the TRP helix (Fig. 5a,b).

**Figure 5.**
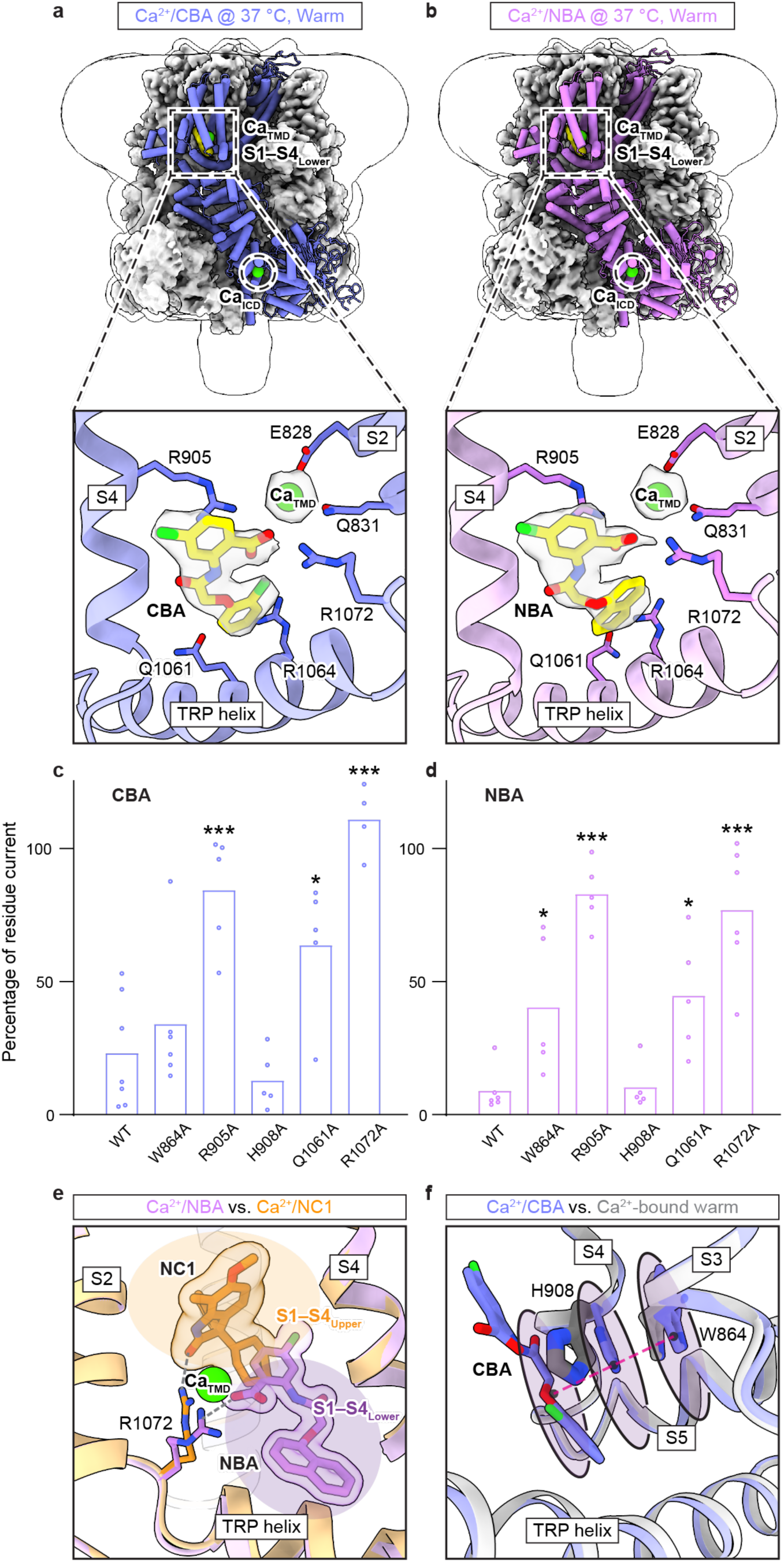
NBA and CBA inhibit TRPM4 through the S1–S4_Lower_ site. **a,b** Structure of TRPM4 in complex with Ca^2+^/CBA at 37 °C (**a**, PDB-ID 9Z23) and Ca^2+^/NBA at 37 °C (**b**, PDB-ID 9Z25). One subunit is shown in cartoon representation, while the remaining three subunits are displayed as the cryo-EM map, viewed parallel to the membrane. The transparent envelope outlines the detergent micelle and the disordered region of the C-terminal coiled-coil. CBA and NBA are highlighted in yellow and Ca^2+^ in green. Bottom panels show the details of the S1–S4 binding site, with the protein backbone in cartoon representation and key interacting residues in stick representation. Densities for CBA and Ca_TMD_ (**a**) and for NBA and Ca_TMD_ (**b**) are shown as transparent surface representations contoured at 17 σ and 20 σ, respectively. The fitted CBA and NBA molecules are shown in stick representation, and Ca_TMD_ as a green sphere. **c,d**, CBA- and NBA-dependent inhibition at +100 mV in cells overexpressing wild-type TRPM4 and binding-site mutants. The percentage of remaining currents after the application of NBA or CBA at 22 °C was calculated by dividing the steady-state current recorded after NBA or CBA application by the steady-state current recorded before application. Each point represents an independently measured cell, with bars indicate the mean. Numbers of independently measured cells are: NBA, 6 (WT), 5 (W864A), 5 (R905A), 5 (H908A), 5 (Q1061A), and 6 (R1072A); CBA, 7 (WT), 6 (W864A), 5 (R905A), 5 (H908A), 5 (Q1061A), and 4 (R1072A). Statistical analysis was performed using one-way ANOVA with Bonferroni’s post hoc test, comparing each mutant with wild type. (*p<0.05, ***p<0.001). The adjusted p-values for each mutant compared to the wild-type (WT) control were as follows: CBA, >0.9999 for W864A, 0.0002 for R905A, >0.9999 for H908A, 0.0149 for Q1061A, and <0.0001 for R1072A; NBA, 0.0458 for W864A, <0.0001 for R905A, >0.9999 for H908A, 0.0179 for Q1061A, and <0.0001 for R1072A. Averaged raw data are shown in Extended Data Fig. 2b. **e**, Structural comparison of the Ca^2+^/NBA-bound state (purple, PDB-ID 9Z26) and the NC1-bound state (orange, PDB-ID 9Z21) of TRPM4, superimposed using S1–S4 (residues 760–910). NBA and NC1 are shown in stick representation with a transparent envelope, and Ca_TMD_ as a green sphere. The filled orange and blue ellipses mark the upper and lower pockets within the S1–S4 domain, respectively. Arg1072 interacts with both ligands, as indicated by the dashed lines. **f**, Structural comparison of the Ca^2+^/CBA-bound state (blue, PDB-ID 9Z23) and the Ca^2+^-bound warm state (grey, PDB-ID 9B8W) of TRPM4, superimposed using S1–S4 (residues 760–910). CBA and the side chains of H908 and W864 are shown as stick representation, forming a triple stacking interaction indicated by the transparent discs and the red dashed line.

Surprisingly, this S1–S4_Lower_ site differs from the NBA binding site previously reported based on lower-resolution cryo-EM maps^41^, which placed the inhibitor ∼10 Å away on the opposite face of the S4 helix (Extended Data Fig. 9c)—within the cleft between the S1–S4 domain and the pore domain (Extended Data Fig. 9d), analogous to the vanilloid pocket in TRPV channels and the NDNA site in TRPM5^42,43^. Careful inspection of both regions in our high-resolution maps revealed no evidence supporting that earlier assignment. Instead, in our NBA- and CBA-bound structures, the vanilloid pocket consistently displayed the same Y-shaped density seen in apo, TPPO-, and NC1-bound structures, indicating a tightly bound endogenous lipid rather than inhibitor occupancy (Extended Data Fig. 9e).

To validate the S1–S4_Lower_ pocket functionally, we introduced mutations at key interacting residues and examined their effects using patch-clamp recordings. Substitution of either R905 or R1072 by alanine completely abolished inhibition (Fig. 5c,d). Moreover, the second aromatic group, a chlorophenyl in CBA or the bulkier naphthyl in NBA, extends downward into a cavity formed between the S4–S5 linker and the TRP helix (Fig. 5a,b). Mutation of Q1061, positioned near this distal aromatic moiety, also markedly reduced inhibition (Fig. 5c,d).

Together, these results establish the S1–S4_Lower_ pocket as the functional binding site for NBA and CBA, defining a second pharmacologically active locus within the S1–S4 domain. This domain thus emerges as a bifunctional pharmacological hub in TRPM4, accommodating both activators (TPPO, NC1, Ca^2+^) and inhibitors (NBA, CBA) through spatially segregated upper and lower sites that converge on shared gating residues such as R1072 on the TRP helix (Fig. 5e). Consistent with this model, mutation of N786—a residue located within the S1–S4_Upper_ region—abolished NC1 activation while leaving NBA-mediated inhibition largely intact (Extended Data Fig. 2c,d). This uncoupling provides functional evidence that the upper and lower S1–S4 pockets are partially independent, despite their close spatial proximity, and can differentially engage ligands to produce distinct pharmacological outcomes.

### NBA and CBA inhibit TRPM4 by locking a pre-open conformation

To elucidate how NBA and CBA inhibit Ca^2+^-dependent activation, we first examined the conformational transitions that underlie canonical TRPM4 gating. In the Ca^2+^-free cold state, W864 on S3 and H908 on S4 form a stabilizing π–π stacking interaction^44–46^. Upon Ca^2+^ binding, coordinated engagement of the Ca_TMD_ and Ca_ICD_ triggers rearrangements in the S1–S4 bundle and TRP helix, repositioning W864 and H908 and thereby disrupting the contact between the intracellular tips of the S3 and S4 helices (Fig. 5f, the gray structure)—a hallmark of Ca^2+^-gated TRPM channel activation^43,47^. This frees the S4–S5 linker, allowing the S5–S6 pore domain to open^10^.

In the Ca^2+^/NBA- and Ca^2+^/CBA-bound structures, this transition is blocked. The distal aromatic moiety of NBA or CBA occupies the position normally adopted by H908 in the activated state, thereby forcing H908 to remain in its apo-like conformation. As a result, H908, W864, and the ligand form a π–π–π triple stacking interaction that restrains the S4–S5 linker and prevents pore opening (Fig. 5f, the purple structure). The overall architecture of the inhibited state closely matches the Ca^2+^-bound warm closed conformation (backbone RMSD 0.7 Å), indicating that NBA and CBA trap TRPM4 in a Ca^2+^-primed but non-conductive pre-open state.

To test this model, we used decavanadate (DVT), a positive modulator that we previously showed binds at the ICD–TMD interface and facilitates the transition of TRPM4 from a Ca^2+^-bound pre-open state to the open state by pulling the pore-lining S6 helix to open the ion-conducting pore^10,48^. Importantly, DVT binding—and the subsequent transition from the pre-open to the open state—is accompanied by an upward tilting of the intracellular domain toward the transmembrane domain^10^. We therefore reasoned that if NBA/CBA stabilize a Ca^2+^-bound pre-open state, they should prevent the conformational changes required for DVT binding and activation, or even preclude DVT binding altogether. To test this prediction, we determined the structure of TRPM4 in the presence of Ca^2+^, DVT, and CBA at 37 °C (Supplementary Fig. 1; Extended Data Fig.10a; Supplementary Table 3). The resulting map contained densities only for Ca^2+^ and CBA, with no evidence of DVT binding, and the protein conformation was identical to the Ca^2+^/CBA-bound closed state (Extended Data Fig. 10b). These data provide direct structural evidence that DVT cannot override NBA/CBA inhibition, supporting our proposed inhibitory mechanism.

Interestingly, this mechanism converges with that of ATP, which also prevents opening by stabilizing the pre-open state, albeit through a distinct intracellular binding site^10^. Unlike ATP, however, NBA and CBA act from the extracellular side and maintain high potency at physiological temperature, offering both accessibility and stability within the native conformational landscape. Thus, NBA and CBA represent promising scaffolds for the design of TRPM4 inhibitors with therapeutic potential.

## Discussion

This study reveals that TRPM4 pharmacology is not static but dynamically shaped by the physiological environment, with temperature and intracellular Ca^2+^ acting as active determinants of ligand binding, efficacy and mechanism. By integrating single-particle cryo-EM and electrophysiology, we reveal that these intrinsic factors cooperatively remodel the conformational landscape of TRPM4, exposing latent binding sites and revealing context-dependent pharmacological behaviors that are obscured under non-physiological conditions.

Physiological context unmasks the true pharmacological profiles of ligands. The small molecule TPPO, long regarded as a TRPM5-specific inhibitor, emerges here as a potent activator of TRPM4, but only when both temperature and intracellular Ca^2+^ reach physiological levels (Fig. 6a). Conversely, NC1, previously characterized as a Ca^2+^-independent agonist, undergoes a functional reversal: at basal Ca^2+^ it activates TRPM4, whereas at elevated Ca^2+^ it becomes functionally antagonized, manifesting as little to no additional effect beyond Ca^2+^ alone, revealing an unexpected inhibitory facet of Ca^2+^ regulation (Fig. 6b). Mechanistically, these contrasting behaviors arise because temperature and Ca^2+^ cooperatively determine the equilibrium between cold and warm conformations of TRPM4, thereby governing access to key residues within the S1–S4 regulatory domain. Thus, ligand efficacy in TRPM4 is not an intrinsic property of the compound alone, but an emergent consequence of its interplay with the conformational and thermodynamic landscape of the channel.

**Figure 6.**
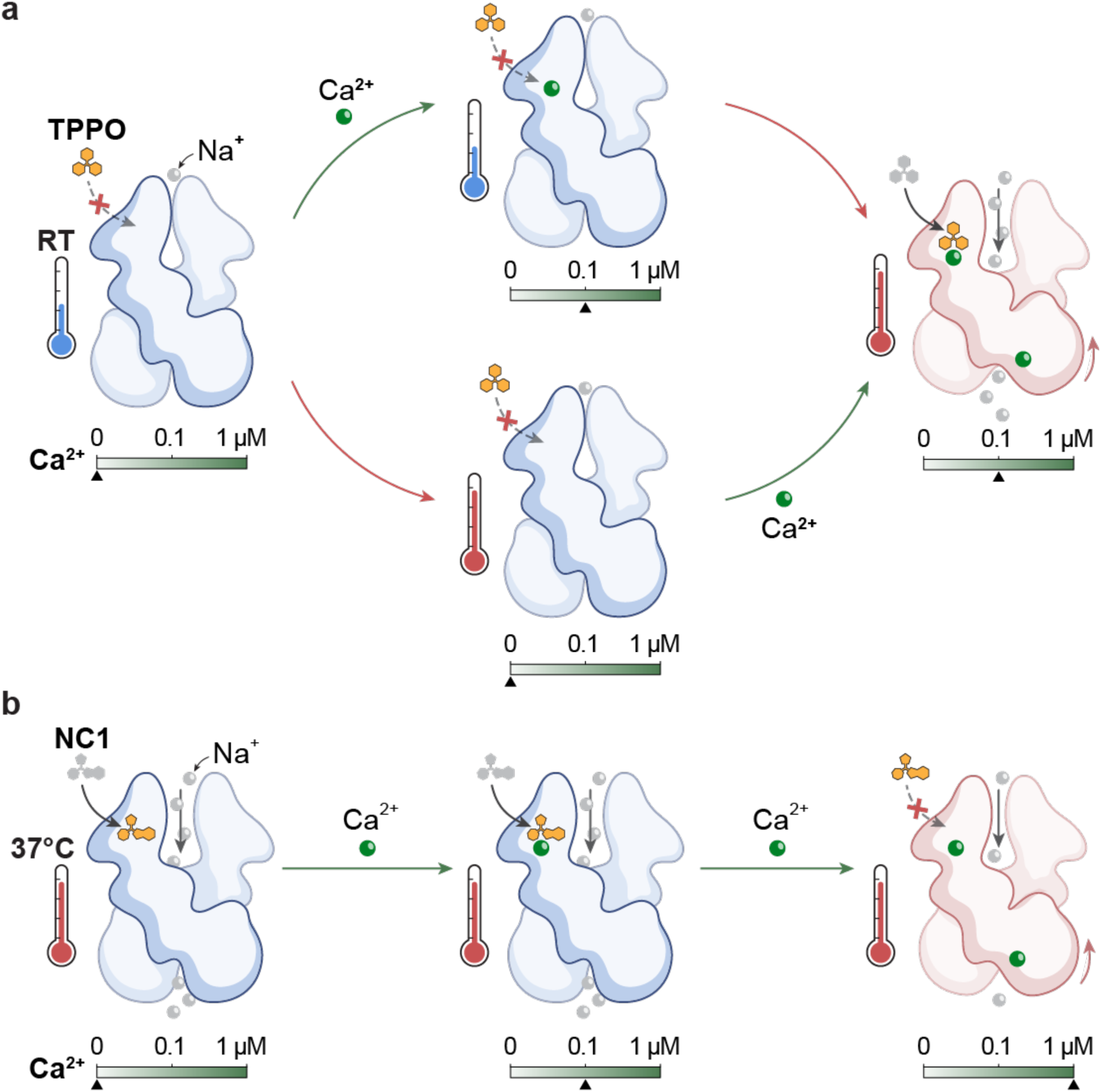
Distinct temperature- and Ca^2+^-dependent activation mechanisms of TPPO and NC1 on TRPM4. **a**, Schematic representation of the polymodal activation of TRPM4 by TPPO, Ca^2+^, and temperature. Neither resting Ca^2+^ levels nor physiological temperature alone is sufficient to enable TPPO-mediated activation. Only when both resting Ca^2+^ and physiological temperature are present can TPPO exert its activating effect. **b**, Schematic representation of NC1 activity, which functions independently of Ca^2+^ at resting levels but loses efficacy once intracellular Ca^2+^ rises to stimulatory concentrations.

Moreover, our structural analyses establish the S1–S4 domain as a versatile pharmacological hub that integrates both environmental and chemical inputs. Within this domain, around the Ca_TMD_ site, two spatially distinct but functionally coupled sites control opposite gating outcomes: the S1–S4_Upper_ pocket, occupied by activators TPPO, NC1, and deacetyl bisacodyl^25^; and the S1–S4_Lower_ pocket, targeted by inhibitors NBA and CBA (Fig. 5e). Both pockets converge on the TRP helix through the conserved residue like R1072 on the TRP helix, which serves as a central allosteric node linking environmental cues to gating transitions. Ligand engagement at the upper pocket promotes activation by coupling between the S1–S4 domain and the pore domain, whereas ligand binding to the lower pocket disrupts this coupling, stabilizing a non-conductive state. This dual-site organization provides a unified explanation for how temperature, Ca^2+^ and chemically diverse ligands—activators and inhibitors alike—modulate TRPM4 through a shared structural framework. Of note, our TPPO- and NC1-bound cryo-EM structures capture TRPM4 in a closed pore conformation, consistent with our previous observations that the pore remains closed in the presence of the endogenous activator Ca^2+^ alone. These observations suggest that agonist engagement with the S1–S4 domain, while necessary for activation, is not sufficient to stabilize an open pore under cryo-EM conditions, likely due to the absence of membrane potential. In contrast, open-pore conformations have been observed in the presence of additional modulators, such as DVT or PIP_2_, which render the channel voltage independent^10,49^.

This regulatory logic uncovered here may represent a general feature of TPRM channels. In TRPM3 and TRPM5, both agonists and antagonists occupy the upper pocket^50–52^; and in TRPM8, ligands engage both^53–55^. The S1–S4 domain thus exemplifies an evolutionary design in which temperature and chemical cues dynamically reconfigure a common structural scaffold to yield distinct, context-dependent functional outcomes.

Notably, we found that temperature-dependent ligand recognition in TRPM4 is not limited to regions that undergo large-scale rearrangements. Even sites within structurally stable domains—such as the S1–S4 transmembrane bundle, which exhibits only subtle temperature-driven movements—can undergo marked shifts in ligand accessibility and efficacy. This principle likely extends to many drug targets, highlighting that seemingly rigid structural regions may harbor cryptic, environment-sensitive pharmacology.

More broadly, these findings support an environment-aware framework of pharmacology, in which physiological variables such as temperature and intracellular Ca^2+^ act as active modulators of ligand binding, potency, and efficacy rather than as passive background conditions. At present, however, the extent to which this principle applies across proteins more generally remains underexplored. Modulation of ligand binding by temperature—and potentially by Ca^2+^ or other physiological cofactors—has historically received limited attention, and relatively few explicit examples are available.

Nevertheless, precedent exists. Crystallographic studies of PBP1B have shown that ligands adopt distinct binding poses when structures are determined at cryogenic versus room temperatures^56^, and analogous temperature-dependent side-chain rearrangements upon metal binding have been observed in hen egg white lysozyme^57^. Similarly, a recent study of ionotropic glutamate receptors has shown that glutamate-dependent pore opening is strongly promoted at physiological temperatures, allowing activated states to be captured by cryo-EM that were not observed in samples prepared under non-physiological temperature conditions^9^. Together with these examples, our results suggest that pharmacological specificity may emerge from the interplay between ligand chemistry and the dynamic, context-dependent structural state of the target protein. We propose that extending similar approaches to other ion channels and proteins will be an important direction for future work, with potential implications for bridging the gap between in vitro pharmacological screening and in vivo drug performance.

For ubiquitously expressed proteins such as TRPM4, these principles carry important therapeutic implications. By exploiting local physiological changes—such as elevated intracellular Ca^2+^ during stress or altered thermal states in disease—future drugs could be tailored to act selectively in pathological contexts while sparing normal function. Such environment-sensitive design strategies may reduce side effects and offer a new route toward precision pharmacology.

## Acknowledgements

We thank for the support with data collection at the Structural Biology Facility at Northwestern University. We appreciate the Structural Biology Facility and Northwestern IT Research Computing and Data Services for computational support. W.L. is supported by National Institutes of Health (NIH) grants (R01HL153219 and R01NS112363, and R35GM138321). J.D. is supported by a McKnight Scholar Award, a Klingenstein-Simon Scholar Award, a Sloan Research Fellowship in neuroscience, a Pew Scholar in the Biomedical Sciences award, and NIH grants (R01NS111031 and R01NS129804). J.H. is supported by American Heart Association postdoctoral fellowship (24POST1196982). A portion of this research was supported by NIH grant R24GM154185 and performed at the Pacific Northwest Center for Cryo-EM (PNCC) with assistance from Marcelo de Farias.

## Author Contributions

J.D. and W.L. supervised the project. J.H. carried out mutagenesis, protein purification, cryo-EM data collection and processing, and structural analysis for all TRPM4 structures included in the study. S.I. and J.C. contributed to the initial cryo-EM studies of TRPM4 in complex with NBA, CBA, and NC1. S.J.P. and J.L. performed functional experiments. G.O. and J.S. carried out mutagenesis and expression test. J.H., J.D. and W.L. wrote the manuscript, with contributions from S.J.P. and J.L. All authors participated in manuscript preparation and approved the final version.

---

**Extended Data Figure 1.**
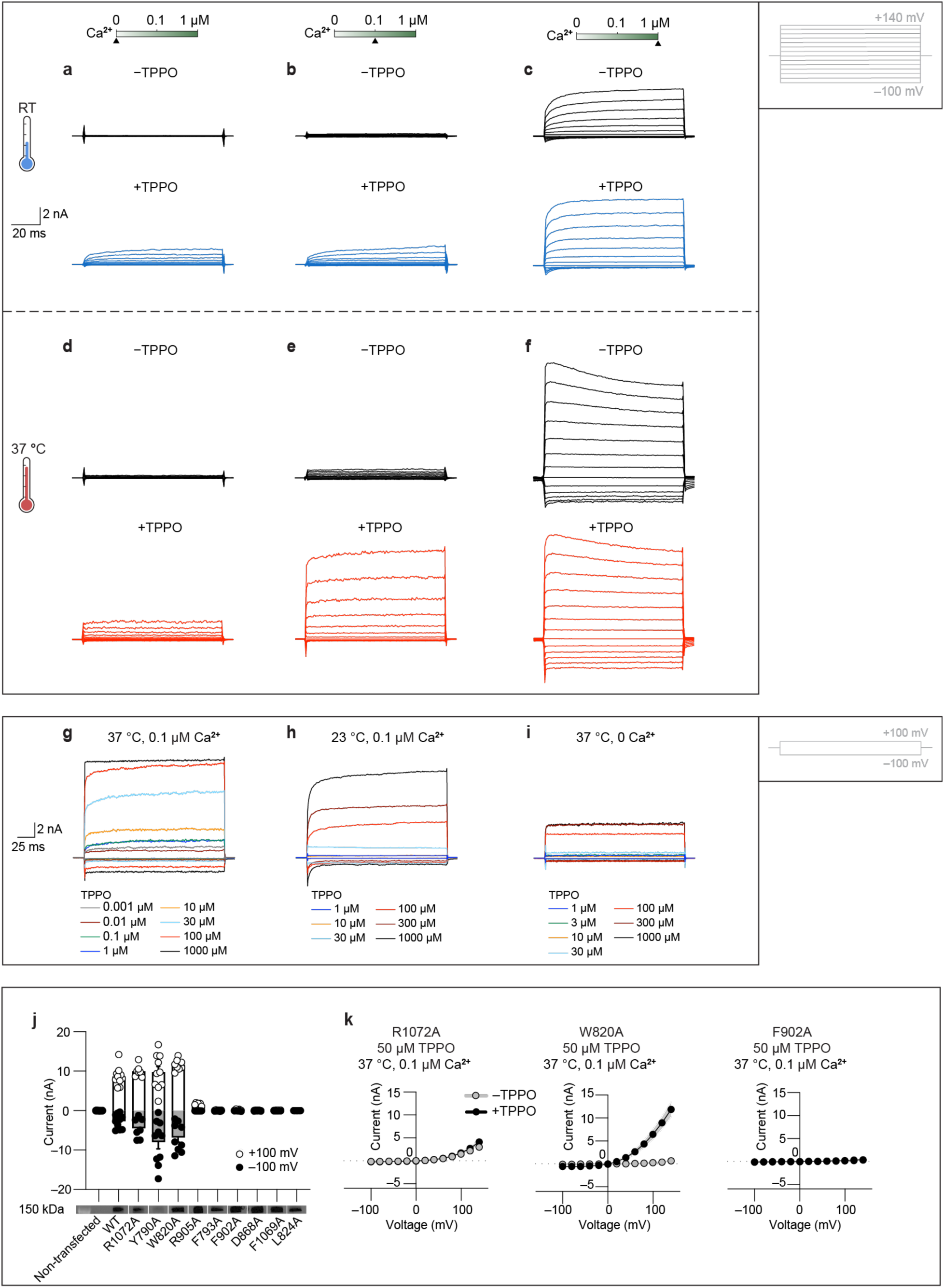
Electrophysiological characterization of TPPO-dependent activation of TRPM4. **a–f**, Representative whole-cell current traces recorded at room temperature (23 °C) and 37 °C with free intracellular Ca^2+^ concentrations of 0 µM (**a,d**), 0.1 µM (**b,e**), and 1 µM (**c,f**), evoked by a voltage-step protocol from –100 mV to +140 mV in 20-mV increments (100 ms per step). The currents without and with 50 µM extracellular TPPO at each condition were measured from a single cell. **g–i**, Representative whole-cell current traces from the TPPO dose response recordings acquired using a two-step protocol (–100 mV followed by +100 mV for 200 ms each) under 0.1 µM Ca^2+^ at 37 °C (**g**), 0.1 µM Ca^2+^ at 23 °C (**h**), and 0 µM Ca^2+^ at 37 °C (**i**). **j**, Whole-cell currents of non-transfected tsA cells and tsA cells overexpressing wild-type TRPM4 and TPPO binding-site mutants evoked by 1 μM free intracellular Ca^2+^ and measured at +100 and −100 mV at room temperature. Each point represents an independently measured cell, with bars indicate the mean. The numbers of independently measured cells are 8 (non-transfected tsA201), 10 (WT), 10 (R1072A), 7 (Y790A), 10 (W820A), 9 (W828A), 12 (S789A), 7 (R905A), 12 (F793A), 6 (F902A), 5 (D868A), 3 (E898A), 4 (F1069A), and 5 (L824A). Cell surface expression of wild-type TRPM4 and mutants is shown, confirming proper trafficking to the plasma membrane; uncropped gels are provided in Supplementary Fig. 2. **K**, Whole-cell currents measured in tsA cells overexpressing the R1072A, W820A and F902A mutants at 37 °C, with 0.1 µM free intracellular Ca^2+^ in the absence or presence of 50 µM extracellular TPPO. A voltage protocol stepping from −100 mV to +140 mV in 20-mV increments was applied, with each voltage step lasting 100 ms. The currents from all measured cells were then averaged and plotted versus voltage to generate the mean I–V curve, with the line representing the mean and the shaded indicating the s.e.m. The numbers of independently measured cells (n) are as follows: R1072A: n=9 (−TPPO) / 10 (+TPPO); W820A: n=10 (−TPPO) / 12 (+TPPO); F902A: n=8 (−TPPO) / 8 (+TPPO)).

**Extended Data Figure 2.**
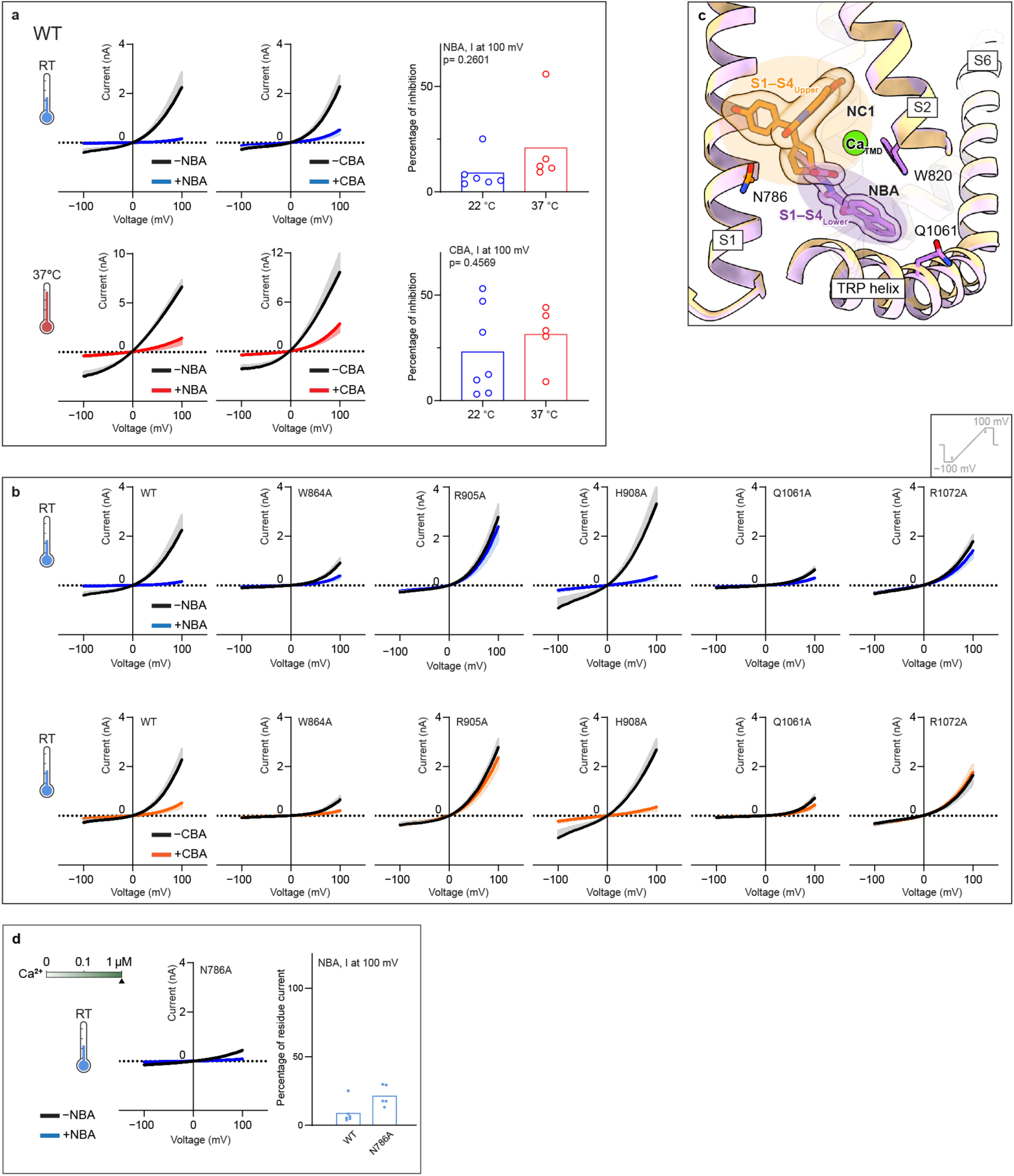
Electrophysiological characterization of TRPM4 inhibition by NBA and CBA. **a**, Left and middle panels show whole-cell currents measured in tsA cells overexpressing wild-type TRPM4 at 22 °C and 37 °C, with 1 µM free intracellular Ca^2+^. A voltage protocol was applied every 5 s to monitor current changes until reaching steady state: −100 mV for 50 ms, ramped to +100 mV over 200 ms, and held at +100 mV for 50 ms. The currents without and with 10 µM extracellular NBA or CBA at each condition were measured from a single cell. The currents from all measured cells were then averaged and plotted versus voltage to generate the mean I–V curve, with the line representing the mean and the shaded indicating the s.e.m. For clarity, the s.e.m. envelope was plotted in one direction. The numbers of independently measured cells are as follows: 6 (NBA, 22 °C), 5 (NBA, 37 °C), 7 (CBA, 22 °C), and 5 (CBA, 37 °C). Right panels show quantification of CBA- and NBA-dependent inhibition at +100 mV for the recordings shown in the left and middle panels. The percentage of remaining currents after the application of NBA or CBA was calculated by dividing the steady-state current recorded after NBA or CBA application by the steady-state current recorded before application. Each point represents an independently measured cell, and bars indicate the mean. A t-test was used to compare inhibition at 22 °C versus 37 °C. A two-sided t-test was performed, with p-values of 0.2601 (NBA) and 0.4569 (CBA). **b**, Whole-cell currents measured in tsA cells overexpressing wild-type TRPM4 or NBA/CBA binding-site mutants at 22 °C, with 1 µM free intracellular Ca^2+^, without and with 10 µM extracellular NBA or CBA. A voltage protocol was applied every 5 s to monitor current changes until reaching steady state: −100 mV for 50 ms, ramped to +100 mV over 200 ms, and held at +100 mV for 50 ms. The currents without and with 10 µM extracellular NBA or CBA at each condition were measured from a single cell. The currents from all measured cells were then averaged and plotted versus voltage to generate the mean I–V curve, with the line representing the mean and the shaded indicating the s.e.m. For clarity, the s.e.m. envelope was plotted in one direction. Measurements and analysis were performed as in panel (**a**). The numbers of independently measured cells (n) are as follows. NBA: n=6 (WT), n=6 (W864A), n=5 (R905A), n=5 (H908A), n=6 (Q1061A), n=6 (R1072A); CBA: n=7 (WT), n=6 (W864A), n=5 (R905A), n=5 (H908A), n=5 (Q1061A), n=4 (R1072A). **c**, Structural comparison of the Ca^2+^/NBA-bound state (purple, PDB-ID 9Z26) and the NC1-bound state (orange, PDB-ID 9Z21) of TRPM4, superimposed using the S1–S4 domain (residues 760–910). NBA and NC1 are shown in stick representation with a transparent envelope, Ca_TMD_ as a green sphere and the protein backbone in cartoon representation and N786 side chain in stick representation. The filled orange and blue ellipses mark the upper and lower pockets within the S1–S4 domain, respectively. **d**, Left panel shows whole-cell currents measured in tsA cells overexpressing the N786A mutant at 22 °C, with 1 µM free intracellular Ca^2+^, without and with 10 µM extracellular NBA in the same cells, averaged from 5 independent cells. Right panel shows quantification of NBA-dependent inhibition at +100 mV for wild-type TRPM4 and the N786A mutant. Measurements and analysis were performed as in panel (**a**). Each point represents an independently measured cell, and bars indicate the mean. The numbers of independently measured cells are 6 (WT), and 5 (N786A). A two-sided t-test was performed, with a p-value of 0.0272.

**Extended Data Figure 3.**
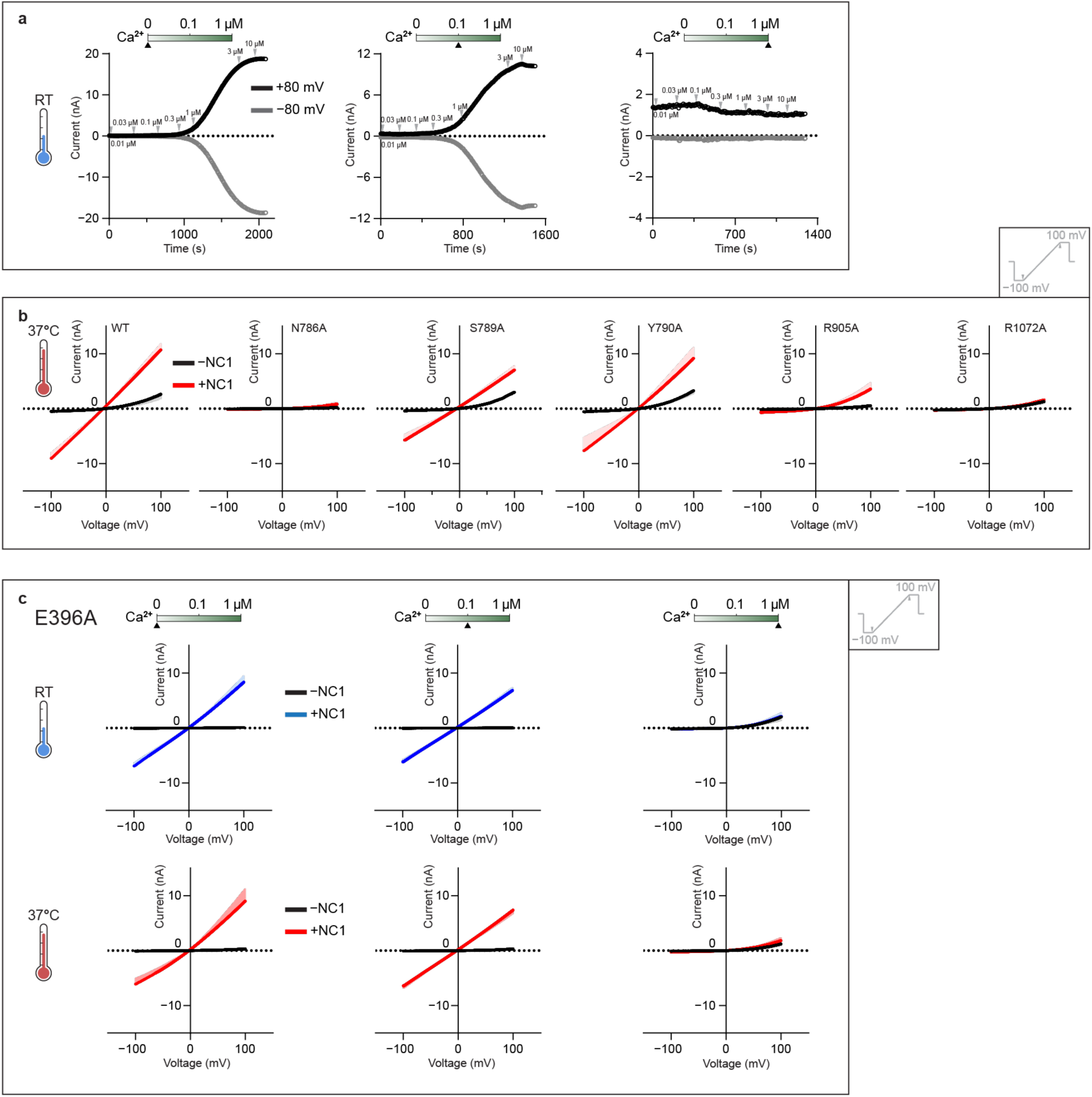
Electrophysiological characterization of NC1-dependent activation of TRPM4. **a**, The representative current traces for the NC1 dose-response measurements for wild-type TRPM4 at 22 °C with free intracellular Ca^2+^ of 0 µM, 0.1 µM, and 1 µM. A voltage protocol was applied every 5 s to monitor current changes: 0 mV for 50 ms, switch to +80 mV for 50 ms, followed by −80 mV for 50 ms, and return to 0 mV for 50 ms. **b**, Whole-cell currents measured in tsA cells overexpressing wild-type TRPM4 or NC1 binding-site mutants at 37 °C, with 0.1 µM free intracellular Ca^2+^, without and with 1 μM extracellular NC1. A voltage protocol was applied every 5 s to monitor current changes until reaching steady state: −100 mV for 50 ms, ramped to +100 mV over 200 ms, and held at +100 mV for 50 ms. The currents without and with NC1 at each condition were measured from a single cell. The currents from all measured cells were then averaged and plotted versus voltage to generate the mean I–V curve, with the line representing the mean and the shaded indicating the s.e.m. For clarity, the s.e.m. envelope was plotted in one direction. The numbers of independently measured cells are 5 (WT), 5 (N786A), 4 (S789A), 5 (Y790A), 5 (R905A), and 4 (R1072A). **c**, Whole-cell currents were measured in tsA cells overexpressing the E396A mutant of TRPM4 at 22 °C (upper panels) and 37 °C (lower panels), with intracellular free Ca^2+^ concentrations of 0 µM, 0.1 µM, and 1 µM. A voltage protocol was applied every 5 s to monitor current changes: −100 mV for 50 ms, ramped to +100 mV over 200 ms and held at +100 mV for 50 ms. The current without or with 1 μM extracellular NC1 in each condition was measured in a single cell. The currents from all measured cells were then averaged and plotted versus voltage to generate the mean I–V curve, with the line representing the mean and the shaded indicating the s.e.m. For clarity, the s.e.m. envelope was plotted in one direction. The numbers of independently measured cells are 5 (0 µM Ca^2+^, 22 °C), 5 (0.1 µM Ca^2+^, 22 °C), 5 (1 µM Ca^2+^, 22 °C), 5 (0 µM Ca^2+^, 37 °C), 5 (0.1 µM Ca^2+^, 37 °C), and 5 (1 µM Ca^2+^, 37 °C).

**Extended Data Figure 4.**
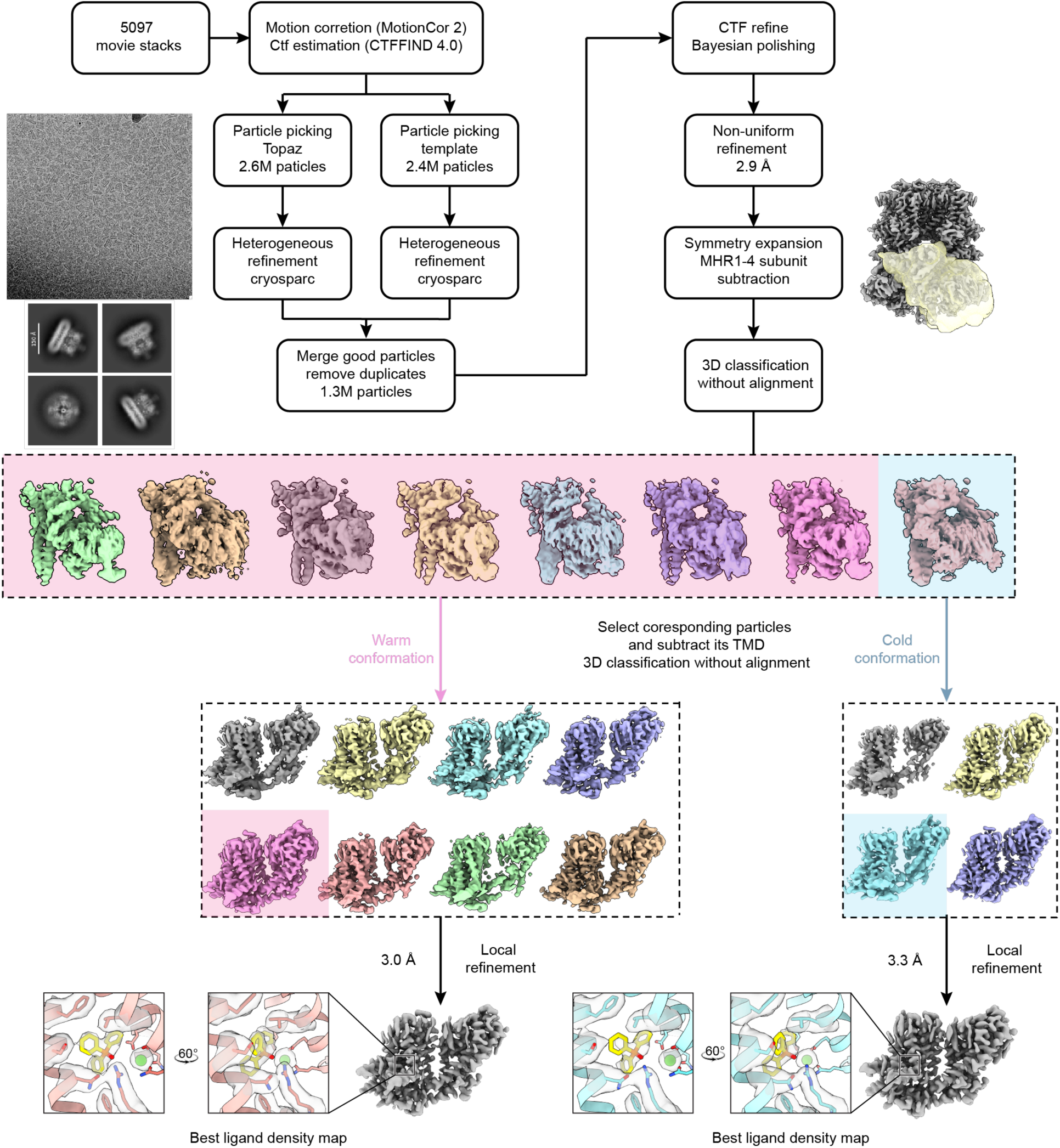
Cryo-EM data processing workflow for the Ca^2+^/TPPO–TRPM4–37 °C dataset. A representative micrograph (out of 5097 micrographs) and representative 2D classification images are shown. The final consensus map and the TMD-focused map with the best-resolved ligand density are displayed with their respective resolutions.

**Extended Data Figure 5.**
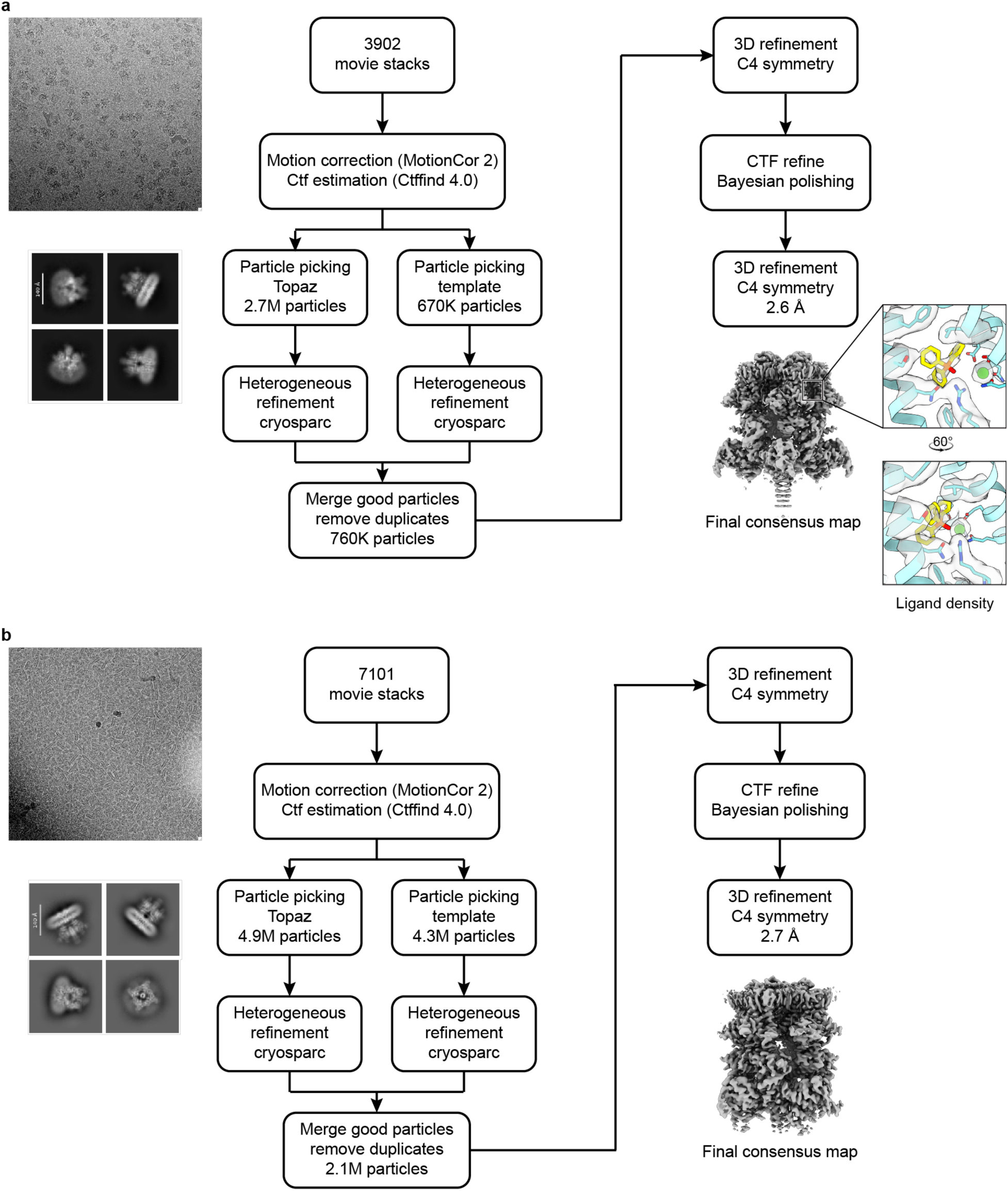
Cryo-EM data processing workflow for the Ca^2+^/TPPO–TRPM4–18 °C (a) and EGTA/TPPO–TRPM4–37 °C (b) datasets. A representative micrograph for Ca^2+^/TPPO–TRPM4–18 °C (**a**, out of 3092 micrographs) and EGTA/TPPO–TRPM4–37 °C (**b**, out of 7101 micrographs), and representative 2D classification images are shown. The final maps are displayed with their respective resolutions.

**Extended Data Figure 6.**
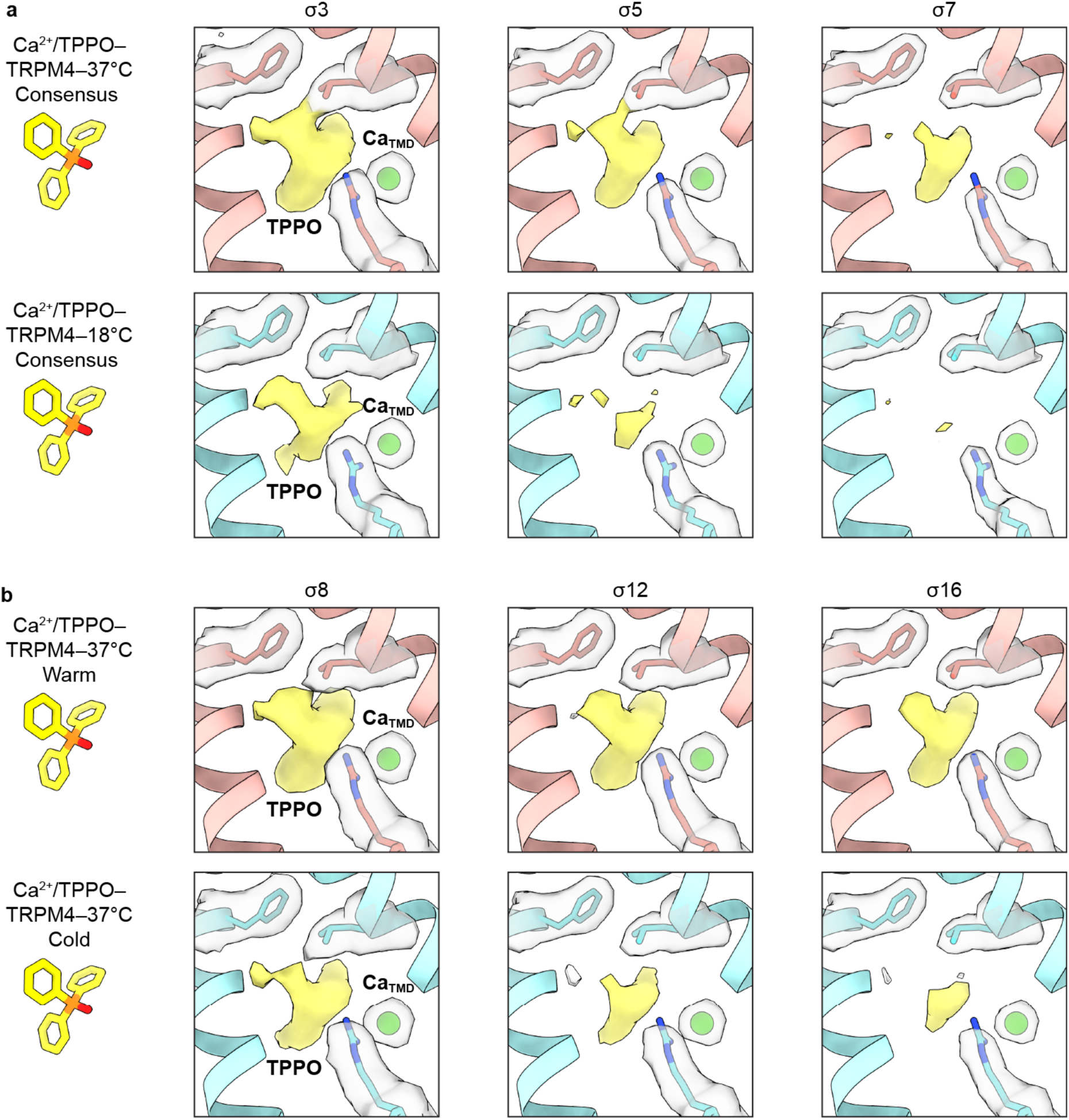
Densities of the TPPO binding site from various datasets or conformations contoured at different σ levels. Densities at the TPPO binding site are shown as yellow surfaces. For comparison, densities for Ca_TMD_ and surrounding interacting residues are displayed as transparent surfaces. The TPPO molecule in the pose fitted to the Ca^2+^/TPPO–TRPM4–37 °C warm map is shown on the left in stick representation.

**Extended Data Figure 7.**
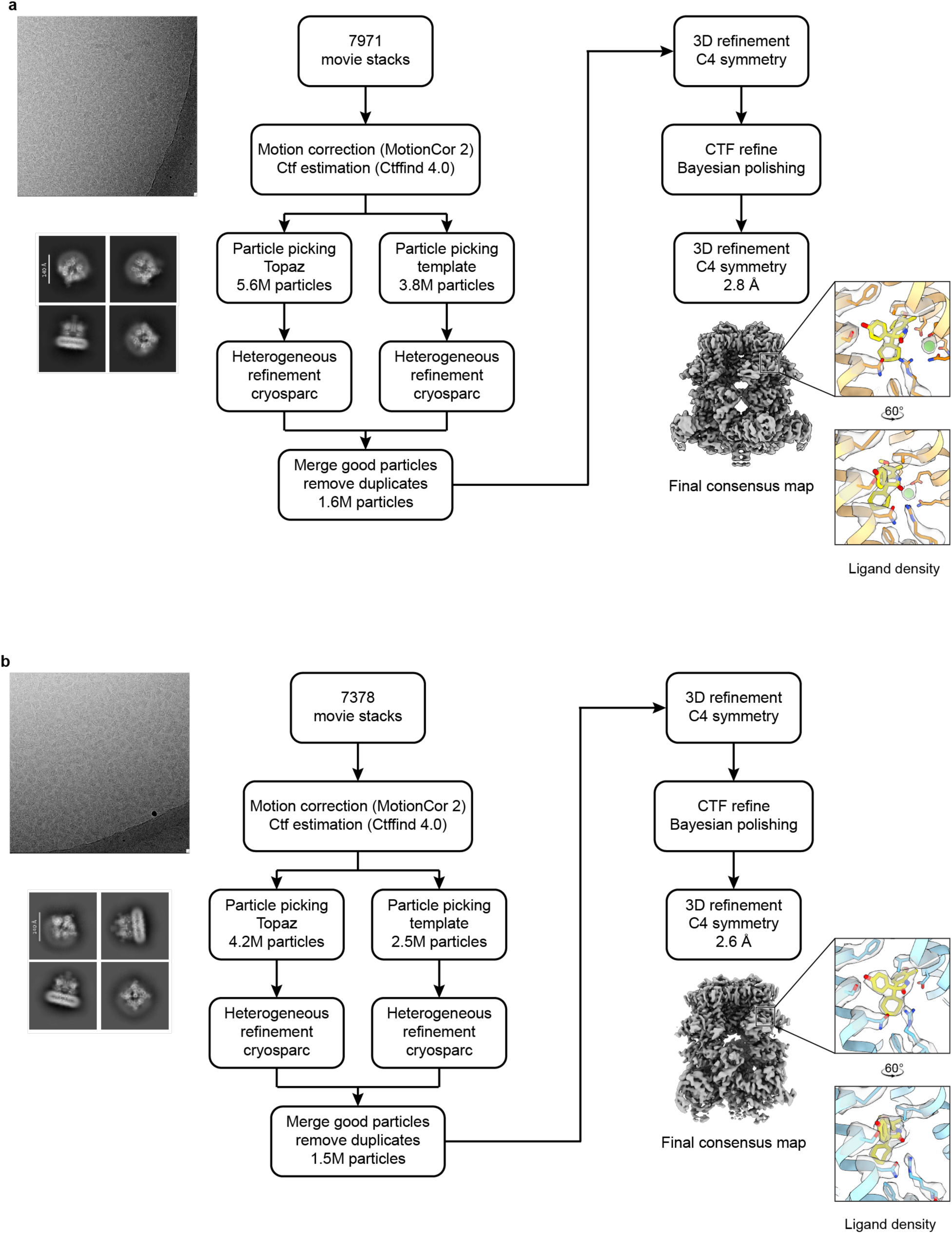
Cryo-EM data processing workflow for the Ca^2+^/NC1–TRPM4–37 °C (a) and EGTA/NC1–TRPM4–37 °C (b) datasets. A representative micrograph for Ca^2+^/NC1–TRPM4–37 °C (**a**, out of 7971 micrographs) and EGTA/NC1–TRPM4–37 °C (**b**, out of 7378 micrographs), and representative 2D classification images are shown. The final maps are displayed with their respective resolutions.

**Extended Data Figure 8.**
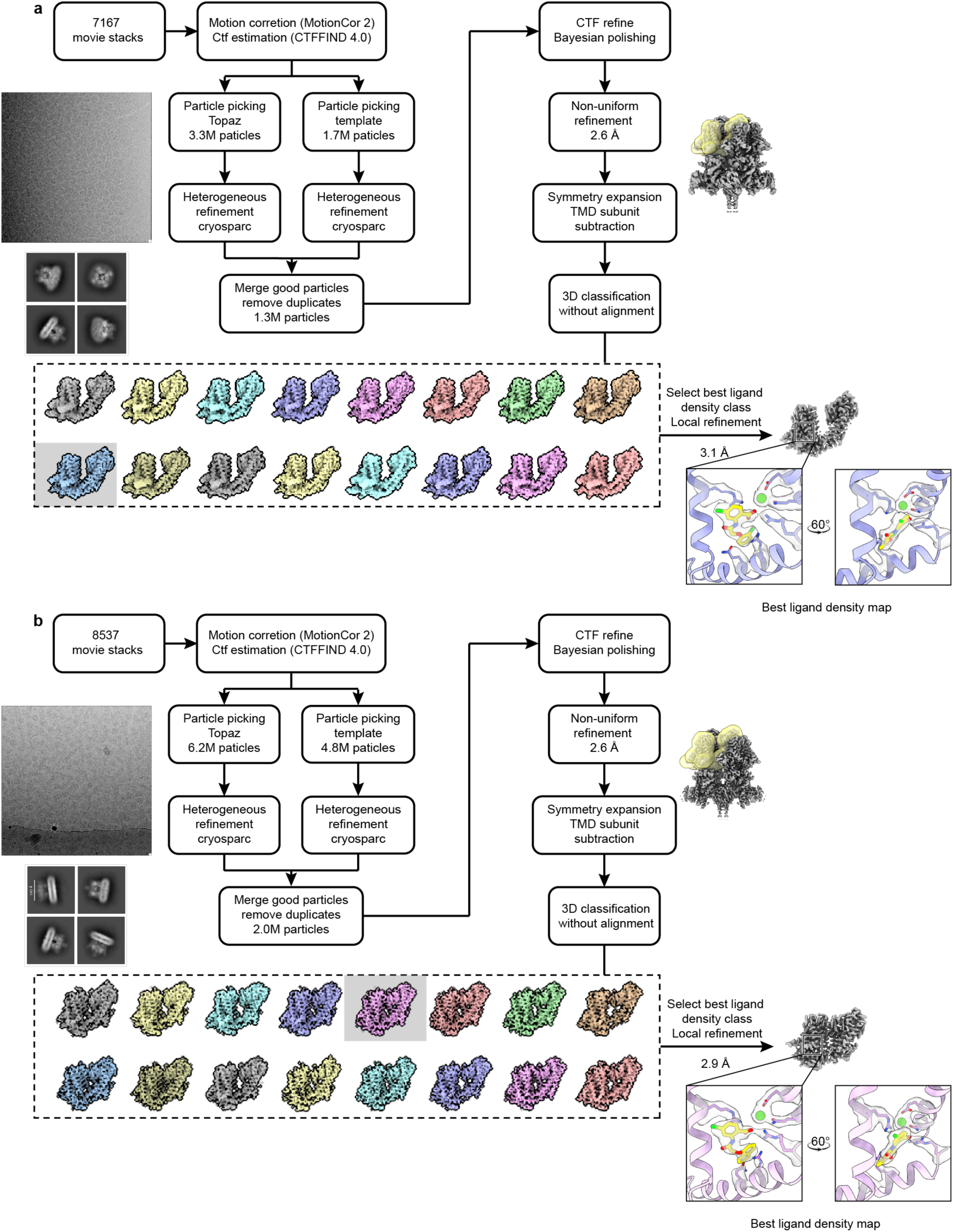
Cryo-EM data processing workflow for the Ca^2+^/NBA–TRPM4–37 °C (a) and Ca^2+^/CBA–TRPM4–37 °C (b) datasets. A representative micrograph for Ca^2+^/NBA–TRPM4–37 °C (**a**, out of 7167 micrographs) and Ca^2+^/CBA–TRPM4–37 °C (**b**, out of 8537 micrographs), and representative 2D classification images are shown. The final consensus map and the TMD-focused map with the best-resolved ligand density are displayed with their respective resolutions.

**Extended Data Figure 9.**
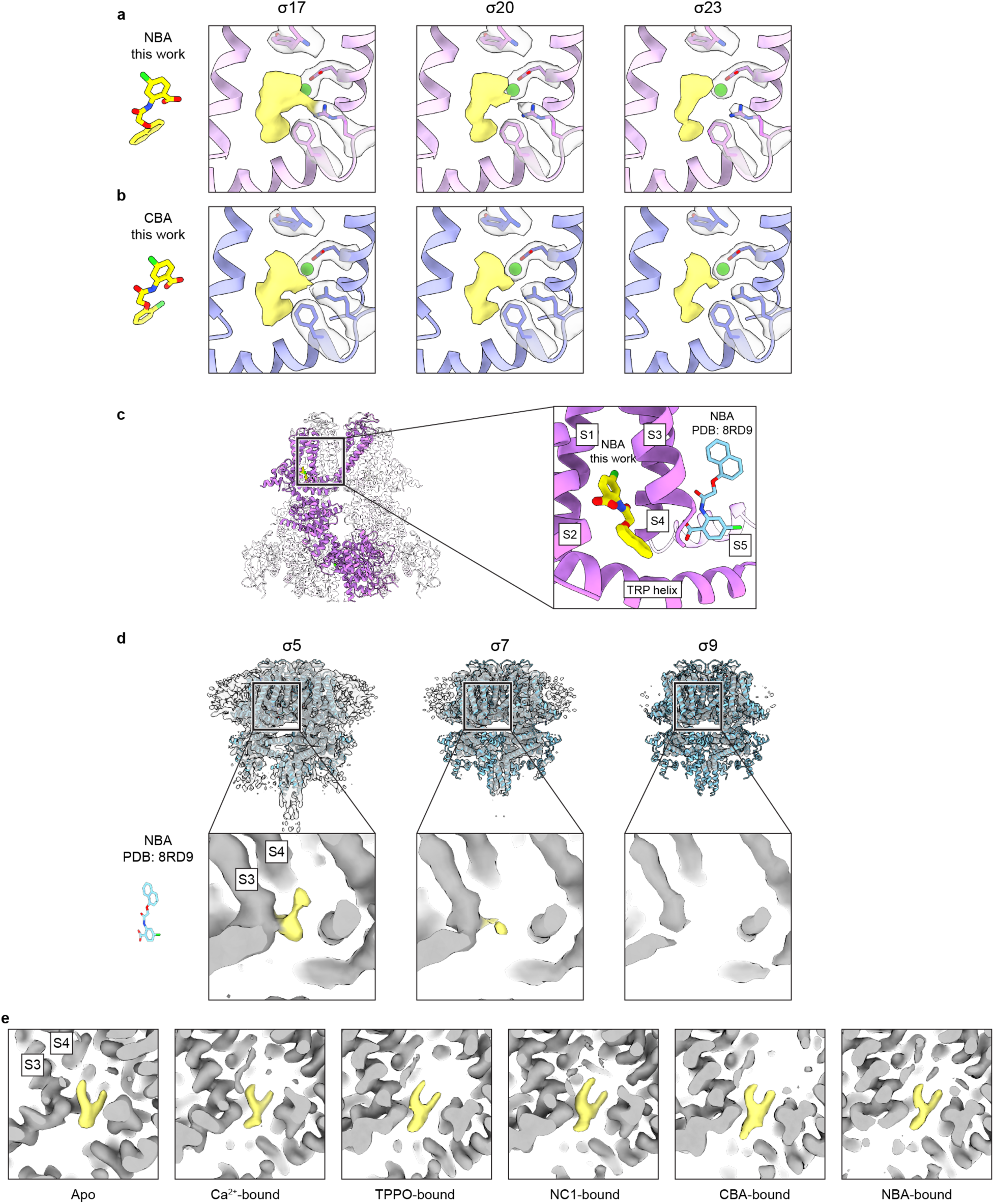
Comparison of the NBA and CBA sites identified in this study with the previously reported NBA site, along with the densities observed within these sites. **a,b**, NBA and CBA densities contoured at different σ levels for the Ca^2+^/NBA–TRPM4–37 °C TMD map and the Ca^2+^/CBA–TRPM4–37 °C TMD map in this study. Densities of NBA and CBA are shown as yellow surfaces. For comparison, densities for Ca_TMD_ and surrounding interacting residues are displayed as transparent surfaces. The NBA and CBA molecules in the poses fitted to the maps are shown on the left in stick representation. **c**, Location of the NBA binding site identified in this work (yellow; PDB-ID 9Z24) compared with the previously reported site (light blue; PDB-ID 8RD9). **d**, The overall cryo-EM map (top) and the densities of the putative NBA binding site (bottom) from the previously published TRPM4 structure in the presence of NBA (EMD-19069), contoured at different σ levels. The putative NBA densities are highlighted in yellow, and the protein densities are shown in gray. The NBA molecule in the pose fitted to the map is shown on the left in stick representation. **e**, Densities at the previously reported NBA binding site in our high-resolution cryo-EM maps across different ligand conditions consistently reveal a Y-shaped density characteristic of a lipid molecule. The Y-shaped densities are highlighted in yellow, and the protein densities are shown in gray.

**Extended Data Figure 10.**
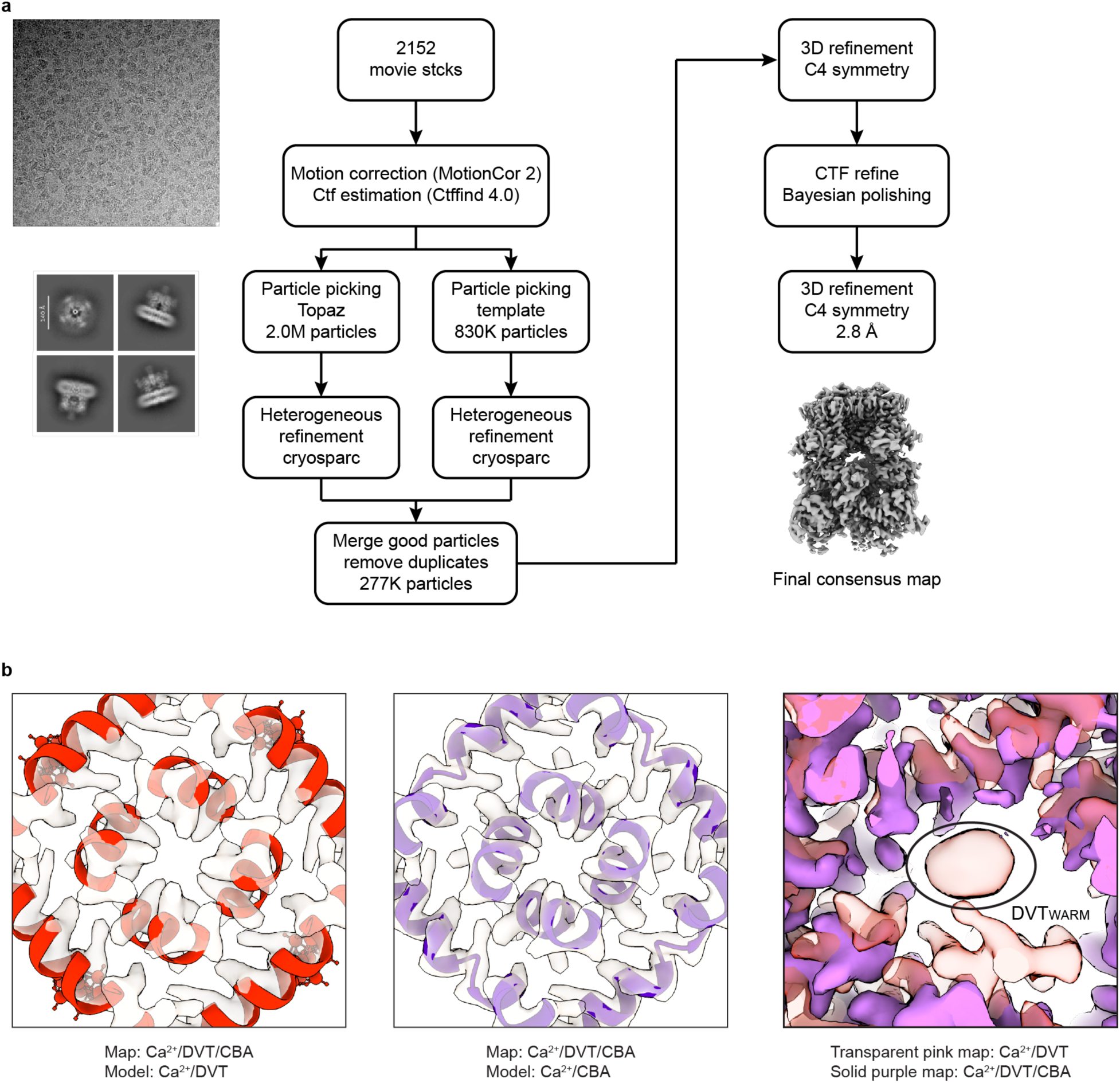
Cryo-EM data processing workflow for the Ca^2+^/DVT/CBA–TRPM4–37 °C dataset. **a**, A representative micrograph (out of 2152 micrographs) and representative 2D classification images are shown. The final map is displayed with the resolution. **b**, Left: superimposition of the Ca^2+^/DVT/CBA–TRPM4 map and the Ca^2+^/DVT–TRPM4_Warm_ model (red; open state), focusing on the transmembrane domain, viewed from the intracellular side. Middle: Superimposition of the Ca^2+^/DVT/CBA–TRPM4 map and the Ca^2+^/CBA–TRPM4 model (purple), focusing on the transmembrane domain, viewed from the intracellular side. Right: superimposition of the cryo-EM maps of Ca^2+^/DVT/–TRPM4_Warm_ (transparent pink) and Ca^2+^/DVT/CBA–TRPM4 (solid purple), focusing on the DVT_Warm_ site and its surroundings.

## Methods

### Human TRPM4 protein expression and purification

The gene encoding human full-length TRPM4 (UniProtKB: Q8TD43) was subcloned into pEG BacMam vector with a 2×Strep tag, GFP and a thrombin-cleavage site at the N terminus as previous described^10,58^. Mutations of TRPM4 were generated use primers provided in Supplementary Data. Bacmid and baculovirus of TRPM4 in a BacMam vector were generated, and P2 viruses were used to infect tsA cells (purchased from the American Type Culture Collection (Catalog CRL-3216) and were not authenticated experimentally in this study) grown in Freestyle 293 expression medium (Gibco) in suspension culture. Cells were incubated at 37 °C for 8 h and10 mM sodium butyrate was added to the culture and the temperature was changed to 30 °C. The cells were collected 72 h after infection and resuspended in a buffer containing 100 mM Tris pH 8.0 and 150 mM NaCl (TBS buffer) in the presence of 1 mM phenylmethylsulphonyl fluoride, 0.8 μM aprotinin, 2 μg ml^−1^ leupeptin and 2 mM pepstatin A. The cells were lysed by sonication and the membrane fraction was collected by centrifugation at 186,000*g* using a 45 Ti rotor (Beckman Coulter) for 1 h at 4 °C. The membrane then homogenized with a Dounce homogenizer in TBS buffer supplemented with protease inhibitors. The protein was extracted from the membrane with TBS buffer supplemented with 1% GDN and protease inhibitors for 3 h at 4 °C. The solubilized proteins were loaded to Strep-Tactin resin.

After washing with TBS buffer supplemented with 0.02% GDN, TRPM4 was eluted with the same buffer, supplemented with 10 mM desthiobiotin. The GFP tag were cleaved, and proteins were concentrated and further purified by size-exclusion chromatography (Superose 6). The peak fractions containing the TRPM4 were pooled and concentrated to 8 mg ml^−1^.

### EM sample preparation and data acquisition

Purified TRPM4 was incubated with 5 mM calcium chloride or 5 mM EGTA, as required for each experiment, for 30 s at 18 or 37 °C. Then 0.1mM TPPO, 0.25mM CBA/NBA or 0.8mM NC1 was added and further incubation for 2 min at 18 or 37 °C respectively. 2.5 μl sample was applied to a glow-discharged Quantifoil holey carbon grid (gold, 2/1 μm size/hole space, 300 mesh). The grids were blotted for 1.5 s in the Vitrobot Mark III set to 100% humidity and 37 °C with a 15 s wait time before being plunge-frozen into liquid ethane cooled by liquid nitrogen.

For the all the sample, images were recorded using the FEI Glacios electron microscope at 200 kV and a nominal magnification of 135,000×. A Falcon 4i direct electron detector with Selectris energy filter was used resulting a pixel size of 0.87 Å or 0.874 Å. EPU was used for automated acquisition. Nominal defocus ranged from −0.5 to −1.4 μm.

### Cryo-electron microscopy data analysis procedure

The detailed workflow for the data-processing procedure is summarized in Extended Data Figs. 4, 5, 7, 8, and 10. In general, the raw movies for each dataset were motion-corrected using MotionCor2 (v.1.1.0)^59^. The per-micrograph defocus values were estimated using ctffind (v.4.1.10)^60^. Particle picking was performed using relion template picker^61^ and topaz (v.0.2.4)^62^. Junk particles were removed by rounds of 3D heterogeneous refinement using CryoSPARC^63^. Good particles were selected for non-uniformed refinement with *C*_4_ symmetry in CryoSPARC to generate a 3D map. Multiple rounds of CTF refinement and Bayesian polishing were performed in RELION and CryoSPARC to further improve the map resolution.

For 37 °C with Ca^2+^ and TPPO dataset, symmetry expansion at the single subunit of ICD level was done from the map refined with *C*_4_ symmetry, followed by monomer ICD subtraction. The subtracted images of the monomer ICD underwent 3D classification without image alignment in RELION to identify the warm and cold conformation. The correspond warm and cold monomer were subsequently undergone similar TMD monomer 3D classification to identify the best the local resolution map with the bound ligand.

For the rest of 37 °C with Ca^2+^ dataset, symmetry expansion at the single subunit of TMD level was done from the map refined with *C*_4_ symmetry, followed by monomer TMD subtraction. The subtracted images of the monomer TMD underwent 3D classification without image alignment in RELION to identify the class with the best local resolution map with the bound ligand. Map resolution estimates were based on the gold standard Fourier shell correlation 0.143 criterion for all datasets.

### Model building

Atomic models were generated by rigid-body fitting of the TMD, MHR1/2 and MHR3/4 and C-terminal domains from a published human TRPM4 model (PDB: 5WP6) into the final cryo-EM maps. Ligands were fitted into the density through real-space refinement using COOT^64^. The CIF file of ligands was generated using grade web server (Global Phasing Ltd.). The models were then manually adjusted in COOT and subjected to phenix.real_space_refine^65^ to improve the model metrics. The final models were validated using phenix.molprobity^66^. Figures were generated using UCSF ChimeraX^67^.

### Electrophysiology

TsA201 cells expressing plasmids encoding N-terminal GFP-tagged human TRPM4 WT and mutants were used. One day after transfection with plasmid DNA (100 ng ml⁻¹) and Lipofectamine 2000 (Invitrogen, 11668019), the cells were trypsinized and replated onto poly-L-lysine-coated (Sigma, P4707) glass coverslips. After cell attachment, the coverslip was transferred to a recording chamber. Whole-cell patch-clamp recordings were performed at room temperature (21–23 °C) or body temperature (36–38 °C). The temperature of perfusion solutions was controlled by thermal control devices (SC-20/CL-100, Warner Instruments).

For whole-cell recordings, glass pipettes were pulled to 3–5 MΩ and filled with an internal solution. Signals were amplified with a MultiClamp 700B amplifier (controlled by MultiClamp Commander v2.2.2.2) and digitized using a Digidata 1550B A/D converter controlled by Clampex v11.3 (Molecular Devices). Whole-cell currents were measured in cells with an access resistance of < 10 MΩ after achieving whole-cell configuration. Whole-cell capacitance was compensated by the amplifier circuitry. Whole-cell currents were recorded using a two-step voltage protocol in which cells were held at –100 mV for 200 ms and then stepped to +100 mV for another 200 ms, repeated every 10 s. An alternative voltage-step protocol was applied, in which cells were stepped from –100 mV to +140 mV in 20-mV increments, with each step lasting 100 ms. A voltage ramp from –100 to +100 mV (200 ms duration) was applied every 5 s from a holding potential of 0 mV. Data were sampled at 10 kHz and low-pass filtered at 2 kHz.

Recordings were analyzed using Clampfit v11.3 (Axon Instruments), GraphPad Prism v10.6.1 (GraphPad Software, Inc., La Jolla, CA), and OriginPro 2024 (OriginLab, Northampton, MA). Data were also analyzed using custom Python scripts, including pyABF for loading and processing ABF electrophysiology files. Electrophysiology files were parsed and analyzed in Python v3.11.8 executed in Spyder v6.0.7 (Anaconda environment) on macOS v26.0.1 (ARM architecture). Data processing used pandas v2.2.2 and numerical operations used NumPy v1.26.4. All custom analysis scripts used in this study are available at https://github.com/junuk861/ephys-analysis-scripts (version v1.0.1) and have been archived on Zenodo (DOI: 10.5281/zenodo.19711152). The bath solution (pH 7.4) contained 150 mM NaCl, 3 mM KCl, 10 mM HEPES, 2 mM CaCl_2_, 1 mM MgCl_2_, and 12 mM mannitol. Patch pipettes were filled with internal solutions containing 150 mM NaCl, 1 mM MgCl_2_, 10 mM HEPES, 5 mM EGTA, and CaCl_2_ adjusted (0 or 1.69 mM) to yield 0 or 0.1 µM free Ca^2+^, respectively (pH 7.3, 22 °C) (source: https://somapp.ucdmc.ucdavis.edu/pharmacology/bers/maxchelator/CaEGTA-TS.htm). For whole-cell recordings of NC1-induced and NBA/CBA-inhibited currents, the extracellular solution contained the following (in mM): 150 NaCl, 10 HEPES and 1 MgCl_2_. The intracellular solution contained (in mM): 150 NaCl, 10 HEPES and 5 EGTA. The intracellular concentration of CaCl_2_ was set to either 0, 1.69 or 4.45 mM (corresponding to 0, 0.1 or 1 µM of free Ca^2+^). The pH was adjusted to 7.4 using NaOH in both the extracellular and intracellular solutions.

TPPO (Sigma-Aldrich, T84603) was prepared daily as a 70 mM stock solution in ethanol and diluted to the desired final concentration in the bath solution immediately before use. NC1 (Necrocide 1, MCE, HY-14307), NBA (4-chloro-2-(2-(naphthalen-1-yloxy)acetamido)benzoic acid, MCE, HY-128172), and CBA (4-chloro-2-[2-(2-chloro-phenoxy)-acetylamino]-benzoic acid, Tocris, 6724) were dissolved in DMSO to a stock concentration of 10, 100, and 50 mM, respectively.

### Surface protein labeling

Surface protein labeling experiments were performed using the Pierce™ Cell Surface Protein Isolation Kit (Thermo Fisher Scientific). Surface and total protein levels of wild-type and mutant TRPM4 were assessed by in-gel fluorescence of the N-terminal GFP tag.

### Statistics & Reproducibility

For electrophysiology, each recording was obtained from a single cell using whole-cell patch clamp, and the recording unit (n) represents the number of cells. Details of experimental design, including whether comparisons were performed between cells or within the same cell and the use of independent cells for each condition, are described in the figure legends and main text.

Experiments were repeated across multiple days using independent cell preparations to account for batch-to-batch variability. Statistical analyses were performed using standard methods as described in the relevant method section and figure legends. Data are presented as mean ± s.e.m. unless otherwise indicated. The number of biologically independent experimental replicates/measurements are indicated in the figure legend.

For cryo-EM, single-particle analysis was used for structure determination. Purified samples were vitrified using standard plunge-freezing methods, and automated data collection and data processing followed established protocols in the field, and detailed descriptions are in methods and extended data figures. Micrograph and particle counts are listed in Supplementary Tables 1–3. Statistical analyses were performed using standard methods as described in the relevant method. All attempts to replicate the structural findings were successful.

No statistical method was used to predetermine sample size. Sample sizes were similar to those generally employed in the field for electrophysiological recordings and cryo-EM imaging. No data were excluded from the electrophysiology analyses. During cryo-EM data processing, particles that clearly did not show target protein-like feature or showed broken/disordered protein domain(s) were excluded, consistent with common practice in the field. The experiments were not randomized. The investigators were not blinded to allocation during experiments and outcome assessment.

## Data Availability

Cryo-EM density maps have been deposited at the EMDB (Electron Microscopy Data Bank) and the Research Collaboratory for Structural Bioinformatics Protein Data Bank (RCS-PDB), respectively. The EMDB accession code and the PDB accession code for Ca^2+^/TPPO–TRPM4–37°C is EMDB-73754 and PDB 9Z1W; Ca^2+^/TPPO–TRPM4–37°C warm TMD is EMDB-73755, PDB 9Z1X; Ca^2+^/TPPO–TRPM4–37°C cold TMD is EMDB-73756, PDB 9Z1Y; Ca^2+^/TPPO–TRPM4–18°C is EMDB-73757, PDB 9Z1Z; EGTA/TPPO–TRPM4– 37°C is EMDB-73758, PDB 9Z20; Ca^2+^/NC1–TRPM4–37°C is EMDB-73759, PDB 9Z21; EGTA/NC1–TRPM4–37°C is EMDB-73760, PDB 9Z22; Ca^2+^/CBA–TRPM4–37°C is EMDB-73761, PDB 9Z23; Ca^2+^/CBA–TRPM4–37°C TMD is EMDB-73762, PDB 9Z24; Ca^2+^/NBA–TRPM4–37°C is EMDB-73763, PDB 9Z25; Ca^2+^/NBA–TRPM4–37°C TMD is EMDB-73764, PDB 9Z26; Ca^2+^/CBA/DVT–TRPM4–37°C is EMDB-73765, PDB 9Z27; respectively. Source data are available with the manuscript online. Data and materials can be obtained from the corresponding authors upon request.

## Code Availability

All custom analysis scripts used in this study are available at https://github.com/junuk861/ephys-analysis-scripts (version v1.0.1) and have been archived on Zenodo (DOI: 10.5281/zenodo.19711152).

---

**Supplementary Table 1:** Cryo-EM data collection, refinement and validation statistics. Data are shown for Ca^2+^/TPPO–TRPM4–37°C, Ca^2+^/TPPO–TRPM4–37°C warm TMD, Ca^2+^/TPPO–TRPM4–37°C cold TMD, Ca^2+^/TPPO–TRPM4–18°C, and EGTA/TPPO–TRPM4–37°C.

|  | Ca<sup>2+</sup>/TPPO–<br>TRPM4–<br>37°C<br>(EMDB-<br>73754)<br>(PDB 9Z1W) | Ca<sup>2+</sup>/TPPO–<br>TRPM4–37°C<br>warm TMD<br>(EMDB-<br>73755)<br>(PDB 9Z1X) | Ca<sup>2+</sup>/TPPO–<br>TRPM4–37°C<br>cold TMD<br>(EMDB-<br>73756)<br>(PDB 9Z1Y) | Ca<sup>2+</sup>/TPPO–<br>TRPM4–18°C<br>(EMDB-<br>73757)<br>(PDB 9Z1Z) | EGTA/TPP<br>O–TRPM4–<br>37°C<br>(EMDB-<br>73758)<br>(PDB 9Z20) |
| --- | --- | --- | --- | --- | --- |
| **Data collection and processing** |  |  |  |  |  |
| Magnification | 135000 | 135000 | 135000 | 135000 | 135000 |
| Voltage (kV) | 200 | 200 | 200 | 200 | 200 |
| Electron exposure<br>(e<sup>−</sup>/Å<sup>2</sup>) | 40 | 40 | 40 | 40 | 40 |
| Defocus range (μm) | -0.5 – -1.4 | -0.5 – -1.4 | -0.5 – -1.4 | -0.5 – -1.4 | -0.5 – -1.4 |
| Pixel size (Å) | 0.87 | 0.87 | 0.87 | 0.87 | 0.874 |
| Symmetry imposed | C4 | C1 | C1 | C1 | C4 |
| Initial particle<br>images (no.) | 2.6M |  |  | 2.7M | 4.9M |
| Final particle images<br>(no.) | 1180K | 882K | 294K | 760K | 1929K |
| Map resolution (Å) | 2.9 | 3.0 | 3.3 | 2.6 | 2.5 |
| FSC threshold | 0.143 | 0.143 | 0.143 | 0.143 | 0.143 |
| Map resolution<br>range (Å) | 2.9 – 246.2 | 3.0 – 246.2 | 3.2 – 246.2 | 2.6 – 246.2 | 2.5 – 246.2 |
| **Refinement** |  |  |  |  |  |
| Initial model used<br>(PDB code) | 5WP6 | 5WP6 | 5WP6 | 5WP6 | 5WP6 |
| Model resolution<br>(Å) | 3.3 | 3.3 | 3.5 | 3.1 | 2.7 |
| FSC threshold | 0.5 | 0.5 | 0.5 | 0.5 | 0.5 |
| Model resolution<br>range (Å) | 3.3 – 246.2 | 3.3 – 246.2 | 3.5 – 246.2 | 3.1 – 246.2 | 2.7 – 246.2 |
| Map sharpening *B*<br>factor (Å<sup>2</sup>) | -120 | -30 | -30 | -100 | -100 |
| Model composition |  |  |  |  |  |
| Non-hydrogen<br>atoms | 30220 | 2912 | 2912 | 29272 | 29312 |
| Protein residues | 3976 | 365 | 365 | 3892 | 3896 |
| Ligands | 12 | 2 | 2 | 12 | 0 |
| *B* factors (Å<sup>2</sup>) |  |  |  |  |  |
| Protein | 133.28 | 83.39 | 82.41 | 123.92 | 103.06 |
| Ligand | 144.31 | 98.78 | 95.88 | 78.23 |  |
| R.m.s. deviations |  |  |  |  |  |
| Bond lengths (Å) | 0.004 | 0.003 | 0.002 | 0.006 | 0.003 |
| Bond angles (°) | 0.469 | 0.448 | 0.419 | 0.493 | 0.452 |
| Validation |  |  |  |  |  |
| MolProbity score | 1.21 | 1.13 | 1.10 | 1.14 | 0.92 |
| Clashscore | 1.89 | 2.41 | 2.24 | 1.69 | 0.97 |
| Poor rotamers (%) | 0.68 | 0 | 0 | 0.5 | 1.13 |
| Ramachandran plot |  |  |  |  |  |
| Favored (%) | 96.15 | 97.49 | 97.49 | 96.63 | 97.58 |
| Allowed (%) | 3.85 | 2.51 | 2.51 | 3.37 | 2.42 |
| Disallowed (%) | 0 | 0 | 0 | 0 | 0 |

**Supplementary Table 2:** Cryo-EM data collection, refinement and validation statistics. Data are shown for Ca^2+^/NC1–TRPM4–37°C, EGTA/NC1–TRPM4–37°C, Ca^2+^/CBA–TRPM4–37°C, Ca^2+^/CBA–TRPM4–37°C TMD and Ca^2+^/NBA–TRPM4–37°C.

|  | Ca<sup>2+</sup>/NC1–<br>TRPM4–<br>37°C<br>(EMDB-<br>73759)<br>(PDB<br>9Z21) | EGTA/NC1–<br>TRPM4–37°C<br>(EMDB-<br>73760)<br>(PDB 9Z22) | Ca<sup>2+</sup>/CBA–<br>TRPM4–37°C<br>(EMDB-<br>73761)<br>(PDB 9Z23) | Ca<sup>2+</sup>/CBA–<br>TRPM4–<br>37°C TMD<br>(EMDB-<br>73762)<br>(PDB 9Z24) | Ca<sup>2+</sup>/NBA–<br>TRPM4–<br>37°C<br>(EMDB-<br>73763)<br>(PDB 9Z25) |
| --- | --- | --- | --- | --- | --- |
| **Data collection and processing** |  |  |  |  |  |
| Magnification | 135000 | 135000 | 135000 | 135000 | 135000 |
| Voltage (kV) | 200 | 200 | 200 | 200 | 200 |
| Electron exposure (e<sup>−</sup>/Å<sup>2</sup>) | 40 | 40 | 40 | 40 | 40 |
| Defocus range (μm) | -0.5 – -1.4 | -0.5 – -1.4 | -0.5 – -1.4 | -0.5 – -1.4 | -0.5 – -1.4 |
| Pixel size (Å) | 0.874 | 0.87 | 0.87 | 0.87 | 0.874 |
| Symmetry imposed | C4 | C4 | C4 | C1 | C4 |
| Initial particle images (no.) | 5.6M | 4.2M | 3.3M |  | 6.2M |
| Final particle images (no.) | 1636K | 1475K | 1290K | 160K | 2049K |
| Map resolution (Å) | 2.8 | 2.6 | 2.6 | 3.1 | 2.6 |
| FSC threshold | 0.143 | 0.143 | 0.143 | 0.143 | 0.143 |
| Map resolution range (Å) | 2.8 – 246.2 | 2.6 – 246.2 | 2.6 – 246.2 | 3.1 – 246.2 | 2.6 – 246.2 |
| **Refinement** |  |  |  |  |  |
| Initial model used (PDB code) | 5WP6 | 5WP6 | 5WP6 | 5WP6 | 5WP6 |
| Model resolution (Å) | 3.1 | 2.9 | 3.2 | 3.3 | 2.9 |
| FSC threshold | 0.5 | 0.5 | 0.5 | 0.5 | 0.5 |
| Model resolution range (Å) | 3.1 – 246.2 | 2.9 – 246.2 | 3.2 – 246.2 | 3.3 – 246.2 | 2.9 – 246.2 |
| Map sharpening *B* factor (Å<sup>2</sup>) | -117 | -105 | -102 | -50 | -106 |
| Model composition |  |  |  |  |  |
| Non-hydrogen atoms | 30300 | 29396 | 30304 | 2740 | 30300 |
| Protein residues | 3976 | 3900 | 3972 | 338 | 3972 |
| Ligands | 12 | 4 | 12 | 2 | 12 |
| *B* factors (Å<sup>2</sup>) |  |  |  |  |  |
| Protein | 107.36 | 100.17 | 108.2 | 101.63 | 114.79 |
| Ligand | 117.04 | 83.25 | 66.02 | 116.92 | 82.15 |
| R.m.s. deviations |  |  |  |  |  |
| Bond lengths (Å) | 0.004 | 0.004 | 0.004 | 0.004 | 0.003 |
| Bond angles (°) | 0.488 | 0.439 | 0.460 | 0.468 | 0.440 |
| Validation |  |  |  |  |  |
| MolProbity score | 1.36 | 1.44 | 1.22 | 1.2 | 1.34 |
| Clashscore | 2.14 | 4.57 | 1.6 | 2.19 | 2.35 |
| Poor rotamers (%) | 0.27 | 0.14 | 1.08 | 0.35 | 0.27 |
| Ramachandran plot |  |  |  |  |  |
| Favored (%) | 94.62 | 96.74 | 95.79 | 96.69 | 95.28 |
| Allowed (%) | 5.38 | 3.26 | 4.21 | 3.31 | 4.72 |
| Disallowed (%) | 0 | 0 | 0 | 0 | 0 |

**Supplementary Table 3:** Cryo-EM data collection, refinement and validation statistics. Data are shown for Ca^2+^/NBA–TRPM4–37°C TMD, and Ca^2+^/DVT/CBA–TRPM4–37°C.

|  | Ca<sup>2+</sup>/NBA–<br>TRPM4–<br>37°C TMD<br>(EMDB-<br>73764)<br>(PDB 9Z26) | Ca<sup>2+</sup>/CBA/DVT–<br>TRPM4–37°C<br>(EMDB-73765)<br>(PDB 9Z27) |
| --- | --- | --- |
| **Data collection and processing** |  |  |
| Magnification | 135000 | 135000 |
| Voltage (kV) | 200 | 200 |
| Electron exposure (e–/Å<sup>2</sup>) | 40 | 40 |
| Defocus range (µm) | -0.5 – -1.4 | -0.5 – -1.4 |
| Pixel size (Å) | 0.874 | 0.87 |
| Symmetry imposed | C1 | C4 |
| Initial particle images (no.) |  | 2M |
| Final particle images (no.) | 239K | 278K |
| Map resolution (Å) | 2.9 | 2.8 |
| FSC threshold | 0.143 | 0.143 |
| Map resolution range (Å) | 2.9 – 246.2 | 3.8 – 246.2 |
| **Refinement** |  |  |
| Initial model used (PDB code) | 5WP6 | 5WP6 |
| Model resolution (Å) | 3.2 | 3.2 |
| FSC threshold | 0.5 | 0.5 |
| Model resolution range (Å) | 3.2 – 246.2 | 3.2 – 246.2 |
| Map sharpening *B* factor (Å<sup>2</sup>) | -50 | -95 |
| Model composition |  |  |
| Non-hydrogen atoms | 2945 | 30292 |
| Protein residues | 364 | 3972 |
| Ligands | 2 | 12 |
| *B* factors (Å<sup>2</sup>) |  |  |
| Protein | 110.32 | 191.28 |
| Ligand | 125.37 | 118.86 |
| R.m.s. deviations |  |  |
| Bond lengths (Å) | 0.004 | 0.002 |
| Bond angles (°) | 0.499 | 0.466 |
| Validation |  |  |
| MolProbity score | 1.11 | 1.34 |
| Clashscore | 2.72 | 3.37 |
| Poor rotamers (%) | 0.97 | 0.54 |
| Ramachandran plot |  |  |
| Favored (%) | 97.77 | 96.64 |
| Allowed (%) | 2.23 | 3.36 |
| Disallowed (%) | 0 | 0 |

**Supplementary Figure 1.**
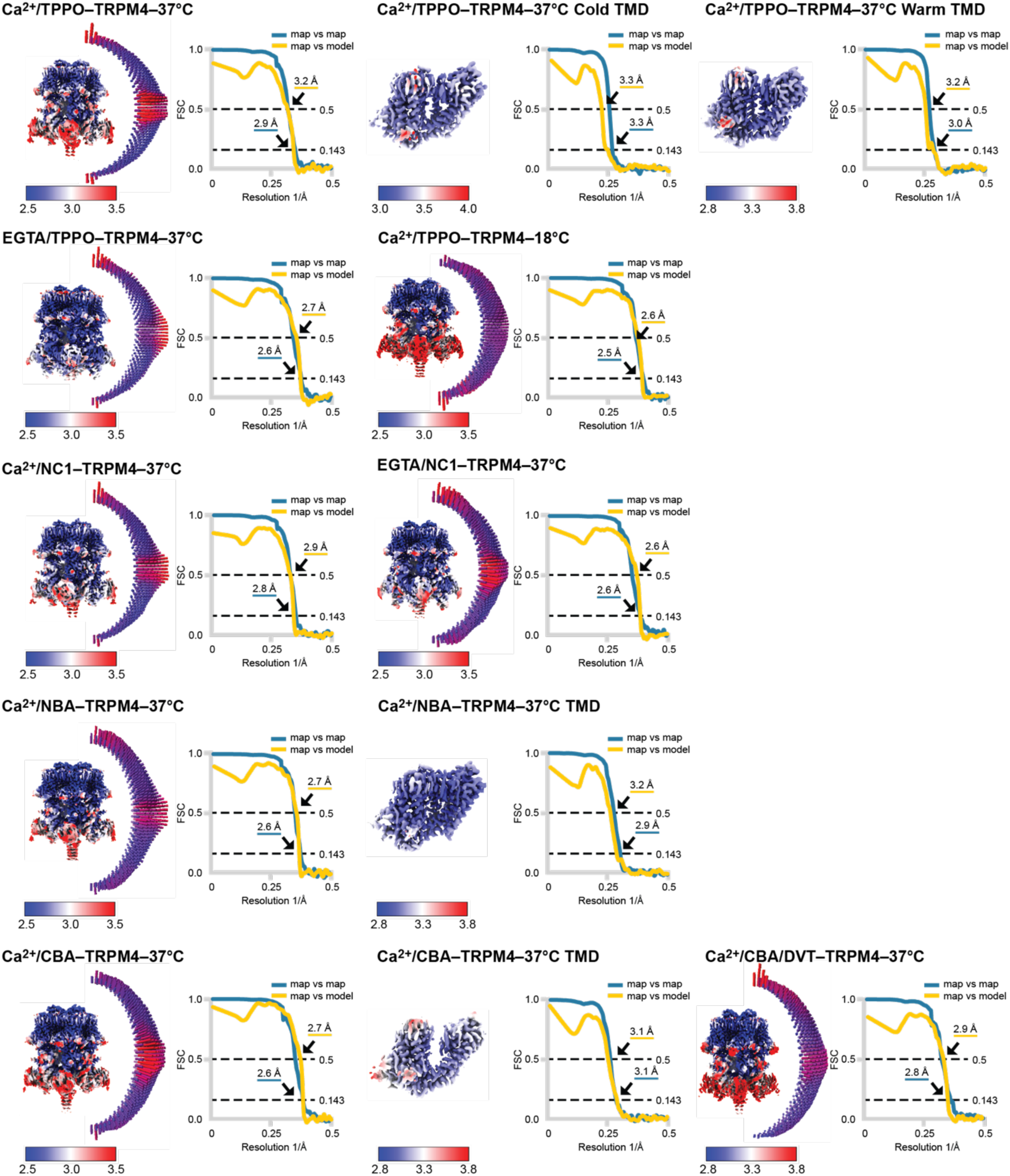
Assessment of cryo-EM data quality. For each map, left: cryo-EM maps colored by local resolution; right: Fourier shell correlation (FSC) plot.

